# BASIN: Background-Anchored Signal Normalization with Integrated Quality Control for CUT&RUN and CUT&Tag

**DOI:** 10.64898/2026.11.21.768281

**Authors:** Harmony R. Salzler, Vasudha Vandadi, Brittany Takushi, Darshan H. Ponkiya, Kami Ahmad, A. Gregory Matera

## Abstract

Quantitative comparison of CUT&RUN and CUT&Tag datasets is complicated when a perturbation changes the global abundance of the profiled chromatin feature or when there is substantial variability between replicates. Read-depth normalization removes global differences by design, whereas exogenous spike-in normalization can introduce substantial technical variability. We developed Background-Anchored SIgnal Normalization (BASIN), an endogenous-reference framework that exploits recurrent, peak-like background in IgG controls as a quantitative internal reference. BASIN Anchor Regions (BARs) act as reproducible read sinks whose representation rises as specific signal falls, allowing global differences to be inferred from the balance between specific and nonspecific reads. BASIN comprises three pipelines: OVERTURE identifies and evaluates BAR panels, HARMONY derives and applies robust sample-specific scale factors from trimmed BAR signal, and CODA determines whether the assumptions supporting normalization are satisfied. BASIN provides a validated, ready-to-use universal BAR panel for *Drosophila melanogaster*, and the OVERTURE pipeline enables BAR panel development in other organisms.

We tested BASIN against independent expectations and compared its performance with spike-in normalization. BASIN remained quantitatively stable across a >100-fold cell-input titration and detected global depletion of H3K36me3 in *Set2* mutants that read-depth normalization concealed. BASIN also preserved expected H3K9me3 differences even when target signal overlapped anchor regions. Applied to a variety of profiling experiments, CODA distinguished normalization failures arising from both biological and technical causes. BASIN thus converts reproducible background from a nuisance signal into an internal reference standard, enabling spike-in-free quantification of global chromatin changes while providing an explicit framework to test when and why that normalization can or cannot be trusted.

## Introduction

Chromatin profiling experiments are interpreted through comparison, and the normalization applied before that comparison determines which differences remain visible. Many perturbations of biological interest change the total abundance of a chromatin feature rather than only its distribution. Loss or inhibition of a histone-modifying enzyme, or mutation of the modified residue itself, can shift signal across much of the genome at once (Meers et al., 2017; Orlando et al., 2014; Penke et al., 2016; Sultana et al., 2026). Sequencing does not directly reflect this, as a sequencing run is of a particular depth, so the read count at a locus reflects that locus’s share of the library rather than its absolute abundance (Chen et al., 2015; Robinson & Oshlack, 2010). When total signal differs between conditions, scaling libraries to equal depth therefore conceals the global difference and can introduce apparent changes of opposite sign at regions that were unaffected (Chen et al., 2015; Orlando et al., 2014). Distinguishing a redistribution of signal from a change in its total amount requires a reference whose behavior is known independently of the feature being measured.

This problem is relevant for CUT&RUN and CUT&Tag, which are now widely used for chromatin profiling in situ (Ahmad & Henikoff, 2025). Both assays produce libraries with low background (Kaya-Okur et al., 2019; Meers et al., 2019b; Skene & Henikoff, 2017), so a large fraction of reads are derived from the target itself. When the target is globally depleted and libraries are nonetheless sequenced to similar depth, the remaining target fragments are sampled more deeply and the depletion is masked. The choice of normalization method to correct for this is consequential: in a recent benchmark of five strategies applied to a single CUT&RUN dataset, fewer than 19% of the peaks called as differentially bound under any method were called under all of them (Ambani et al., 2026).

Existing strategies fall into four classes, each of which resolves the problem in a way that is vulnerable to specific failure modes (Fig. 1A) (Ambani et al., 2026; Patel et al., 2024). Read-depth methods (CPM, RPGC) scale each library to its own sequencing depth and require no exogenous material (Ramírez et al., 2016). However, they enforce equal total signal across samples by design, so a genome-wide change in abundance is removed rather than measured (Chen et al., 2015; Orlando et al., 2014). Distribution-based methods, including quantile transformation and *z*-scoring, likewise rely only on internal signal, but alter the shape of the signal distribution itself (Bolstad et al., 2003; Cheadle et al., 2003). Relative structure within a sample is retained while absolute levels cannot be compared between samples (Bonhoure et al., 2014; Hicks & Irizarry, 2015). Related approaches instead derive a single scale factor from a feature of the signal distribution, such as the slope of the cumulative read distribution (ChIPseqSpikeInFree; Jin et al., 2020) or the 99th percentile of local signal maxima (GNOMES; Roule & Akizu, 2026). These require no exogenous material, but take as their reference a quantity that is itself drawn in part from target signal.

**Figure 1.**
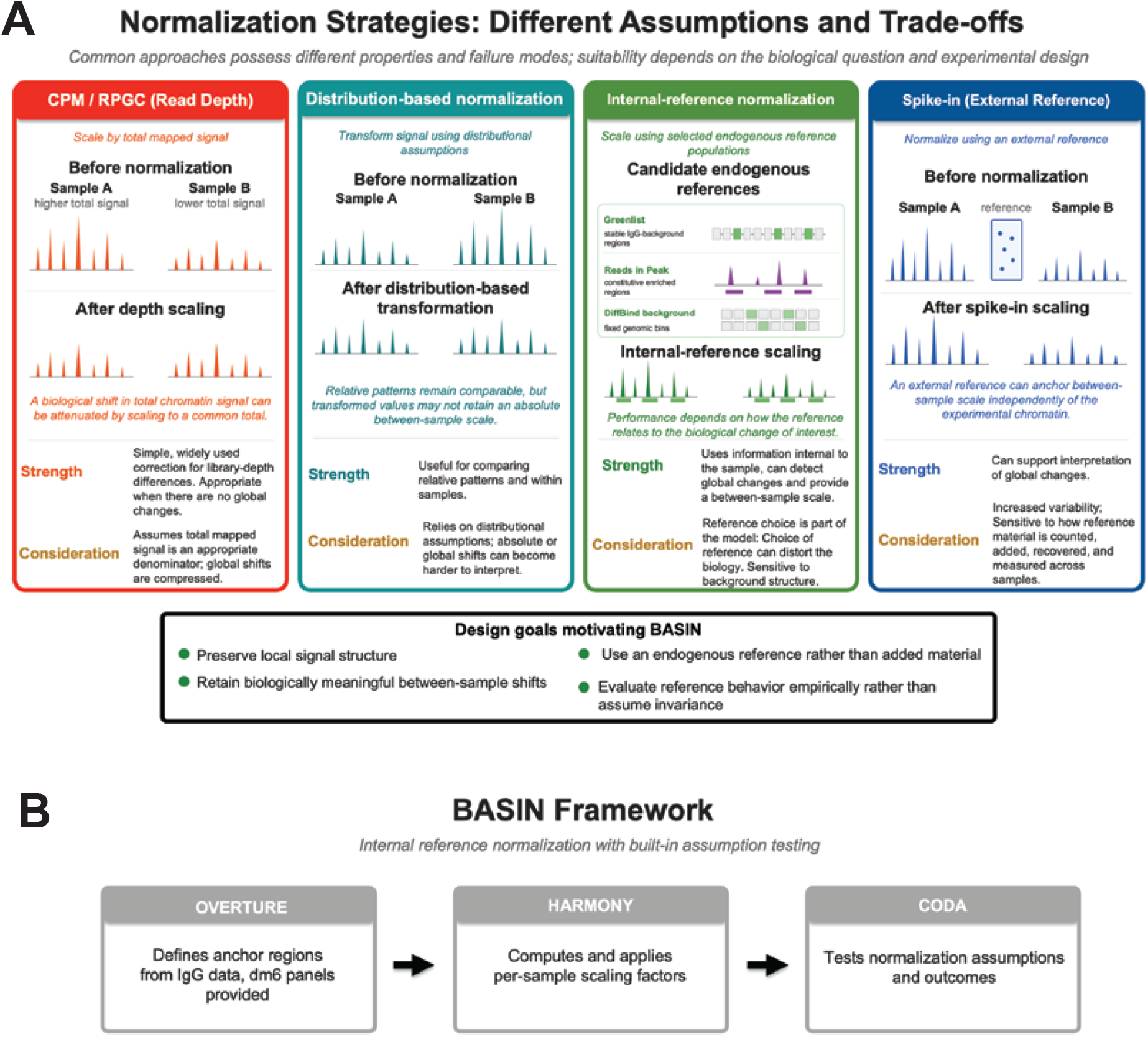
No single existing normalization strategy satisfies all desirable properties for chromatin profiling data. **(A)** Schematic comparison of read-depth (CPM/RPGC), distribution-based, internal-reference, and spike-in normalization, illustrated with idealized coverage. For endogenous internal-reference normalization, three choices of reference are shown: Greenlist regions, consensus peaks (reads in peaks), and DiffBind background bins. Strengths and considerations are listed for each category; the lower box lists properties for an ideal normalization that served as design goals for BASIN. **(B)** The BASIN framework comprises three pipelines. OVERTURE defines BASIN Anchor Regions (BARs) from IgG datasets; a universal *Drosophila melanogaster* dm6 panel is provided with the software, and users may also generate universal panels for other organisms, or experiment-specific panels. HARMONY computes a per-sample scale factor from BAR signal and applies it to generate normalized coverage tracks. CODA evaluates whether the assumptions underlying BASIN normalization were satisfied in a given experiment and returns sample– and experiment-level quality recommendations.

Spike-in normalization addresses the global problem directly, scaling each sample by reads derived from an externally-derived reference (Bonhoure et al., 2014; Orlando et al., 2014). In principle this preserves both local structure and true global differences. In practice, its accuracy depends on cell counting, reproducible addition of reference material, and comparable recovery of reference and experimental chromatin (Patel et al., 2024). Each is a source of technical error and variability, and the contribution of this technical error is not reversible after sequencing. Additionally, spike-in approaches that control for antibody binding efficiency are limited to targets that are shared between the two species, and if the two species are closely related, this can result in a significant proportion of reads discarded due to multi-mapping (Orlando et al., 2014; Patel et al., 2026). Consistent with these concerns, a recent benchmark found spike-in to be the least reliable of five normalization strategies for CUT&RUN (Ambani et al., 2026).

A fourth class has recently emerged as a promising alternative to read-depth, spike-in, and distribution-based normalization. Background based approaches scale to an endogenous reference: a subset of the sample’s own signal presumed to be independent of the biological perturbation under study (Liang & Keleş, 2012). DiffBind (Ross-Innes et al., 2012; Stark & Brown, 2011), csaw (Lun & Smyth, 2016), and Greenlist (de Mello et al., 2024) normalization frameworks all adopt this strategy. These methods require no exogenous material and retain the capacity to preserve global changes. Therefore, they are not subject to the technical failure modes considered above, instead performance is largely determined by the appropriateness of the reference regions. A reference that contains, or responds to, the signal being measured will partially normalize away the effect it was intended to quantify.

What can serve as such a reference depends on what constitutes background in these assays. In CUT&RUN and CUT&Tag, the nonspecific fraction can be considered as two conceptually separable components: (1) cleavage directed by nonspecific antibody binding and (2) nonspecific cutting by MNase in CUT&RUN or Tn5 transposase in CUT&Tag (Kaya-Okur et al., 2019; Meers et al., 2019b; Skene & Henikoff, 2017). Nonspecific cutting can itself be accessibility-driven and peak-like (Henikoff et al., 2020; Hu et al., 2026; Meers et al., 2019a; Thompson & Byrd, 2025) or relatively stochastic, producing a lower-level baseline (Kaya-Okur et al., 2019; Meers et al., 2019b; Skene & Henikoff, 2017). Background signal in sequencing-based chromatin assays is known to exhibit reproducible genomic structure related to features such as mappability and local chromatin properties rather than being uniformly distributed across the genome (Nordin et al., 2023; Rozowsky et al., 2009). The structure and relative contribution of background can also vary with assay chemistry, as CUT&RUN, CUT&Tag, and ChIP-seq differ in their signal and background architecture (Henikoff et al., 2020; Nordin et al., 2023; Oh et al., 2026; Thompson & Byrd, 2025). Strong, reproducible nonspecific signal in heterochromatic and blacklist-associated regions has been noted before, but has generally been filtered rather than exploited (Jain et al., 2015; Teytelman et al., 2013).

We reasoned that endogenous normalization is most promising because it is less susceptible to technical artifacts and can be performed retroactively, but that the choice of normalization regions could potentially be improved. Hence, this study describes Background-Anchored Signal Normalization (BASIN), an internal-reference based method in which reference regions are defined empirically from read sinks that recur in IgG controls, selected for recurrence across assays and biological contexts, and high dynamic range. High signal reference regions likely containing true signal are subsequently trimmed from the reference pool. BASIN also includes an explicit QC framework that evaluates whether its assumptions were met in a given experiment, which to our knowledge other CUT&RUN and CUT&Tag normalization workflows do not provide comprehensively.

BASIN is implemented as three pipelines (Fig1B). OVERTURE allows independent users to create their own organism-specific or experiment-specific reference panels. Here, we demonstrate the generation and utility of *Drosophila melanogaster* specific panels and provide these reference files to the user. In cases where BASIN Anchor Region (BAR) panels have been constructed and/or shared, OVERTURE is unnecessary. HARMONY normalizes experimental samples using a scaling factor generated from signal in samples over BARs using a trimmed-median approach. Finally, CODA evaluates whether BASIN’s core assumptions were satisfied, identifies specific failure modes, and assesses whether normalized output files are satisfactory at both the sample and experimental level.

We evaluated BASIN against independent expectations rather than against agreement with another normalization method. In *Drosophila* mutants expected to lose H3K36me3 genome-wide, BASIN recovered depletion of signal in Set2Δ that was not detectable after read-depth normalization. Across a cell-input titration in which chromatin state was held constant, BASIN-normalized signal remained independent of input, as predicted. In an H3K9me3 dataset in which anchor regions and target signal overlap, upper-tail trimming preserved the expected genotype effect. Applied across a variety of experiments, CODA separated normalization failures with distinct causes. Together, these analyses indicate that peak-like IgG background can serve as an internal reference for quantitative comparison of CUT&RUN and CUT&Tag data, and that the assumptions upon which it rests can be examined in each experiment.

## Results

### BASIN scales each sample to its own background at IgG-defined read sinks

The central premise of BASIN, shared with other endogenous-reference normalization approaches, is that specific and non-specific signal compete for representation within a sequencing experiment (Chen et al., 2015; Orlando et al., 2014). Unlike background-based normalization strategies that preferentially use low-level baseline signal, BASIN exploits regions that recurrently accumulate peak-like signal in IgG controls and uses their collective behavior to estimate the background component of each experimental library. Regardless of their mechanistic origin, BARs serve as “read sinks” that compete for representation with regions of specific signal.

This choice of reference population differs from other endogenous normalization strategies in important ways. Greenlist, the closest conceptual relative to BASIN, identifies highly stable background regions using Shannon-entropy-based criteria and uses these as endogenous normalization factors (de Mello et al., 2024). Although Greenlist also selects regions from a diverse set of IgG controls, it selects for the “most stable” regions, whereas BASIN explicitly selects for regions with a dynamic range capable of revealing titration-like behavior. Peak calling on IgG controls and consensus filtering identifies reproducible, relatively high-count background sinks with substantial dynamic range whose abundance can vary coherently with the background fraction of an experimental library. Other methods rely, implicitly or explicitly, on interpeak background (Diaz et al., 2012; Liang & Keleş, 2012; Lun & Smyth, 2016; Stark & Brown, 2011), but in CUT&RUN and CUT&Tag this diffuse signal is sparse and can vary with reaction conditions (Kaya-Okur et al., 2019; Meers et al., 2019a; Meers et al., 2019b). We reasoned that recurrent IgG-associated signal would be a more informative reference, because it accumulates at defined regions and its share of reads in an antibody-targeted library rises as specific signal falls.

Another deliberate choice is which IgG controls are used for BAR discovery. Universal BAR panels are derived from IgG peaks consistently present across public and in-house datasets present in both assay types (CUT&RUN and CUT&Tag) and diverse biological sources (cell types, tissues, and developmental stages). Requiring presence in both assay types is designed to enrich for background that is robust to assay chemistry, while minimizing background specific to one enzyme (Thompson & Byrd, 2025). Requiring presence across biological sources is likewise intended to limit accessibility peaks that reflect cell-type-specific function. Greenlist, by contrast, generates separate normalization region lists for CUT&RUN and CUT&Tag (de Mello et al., 2024).

The HARMONY pipeline is designed to impute robustness at multiple steps through ranking and trimming signal intervals on a per-sample basis. To compute a scaling factor for normalization (Fig. 2), HARMONY summarizes mean coverage at every BAR from raw, library-size-uncorrected bigWig files at 1-bp resolution. These raw values differ between samples because of both variation in sequencing depth and variation in the relative abundance of nonspecific background. Mean BAR signal values are ranked in descending order within each sample, and the upper fraction – the anchor “drop” threshold, comprising BARs most likely to contain true antibody-associated enrichment rather than background – is removed per sample, so that sample– or genotype-specific variation at individual loci does not influence the scaling factor. The drop threshold default is 20%, but fully is configurable based on target abundance. BARs showing zero signal across all samples are also removed. The median of the retained background-representative values (medBAR, reported as median_kept) defines the HARMONY coverage multiplier as 1/medBAR. BASIN normalization can also be incorporated into common differential-binding frameworks, including DESeq2, edgeR, and DiffBind, by supplying the medBAR-derived sample normalization information through their size-factor or offset interfaces (see software documentation for details) (Love et al., 2014; Robinson et al., 2010; Stark & Brown, 2011).

**Figure 2.**
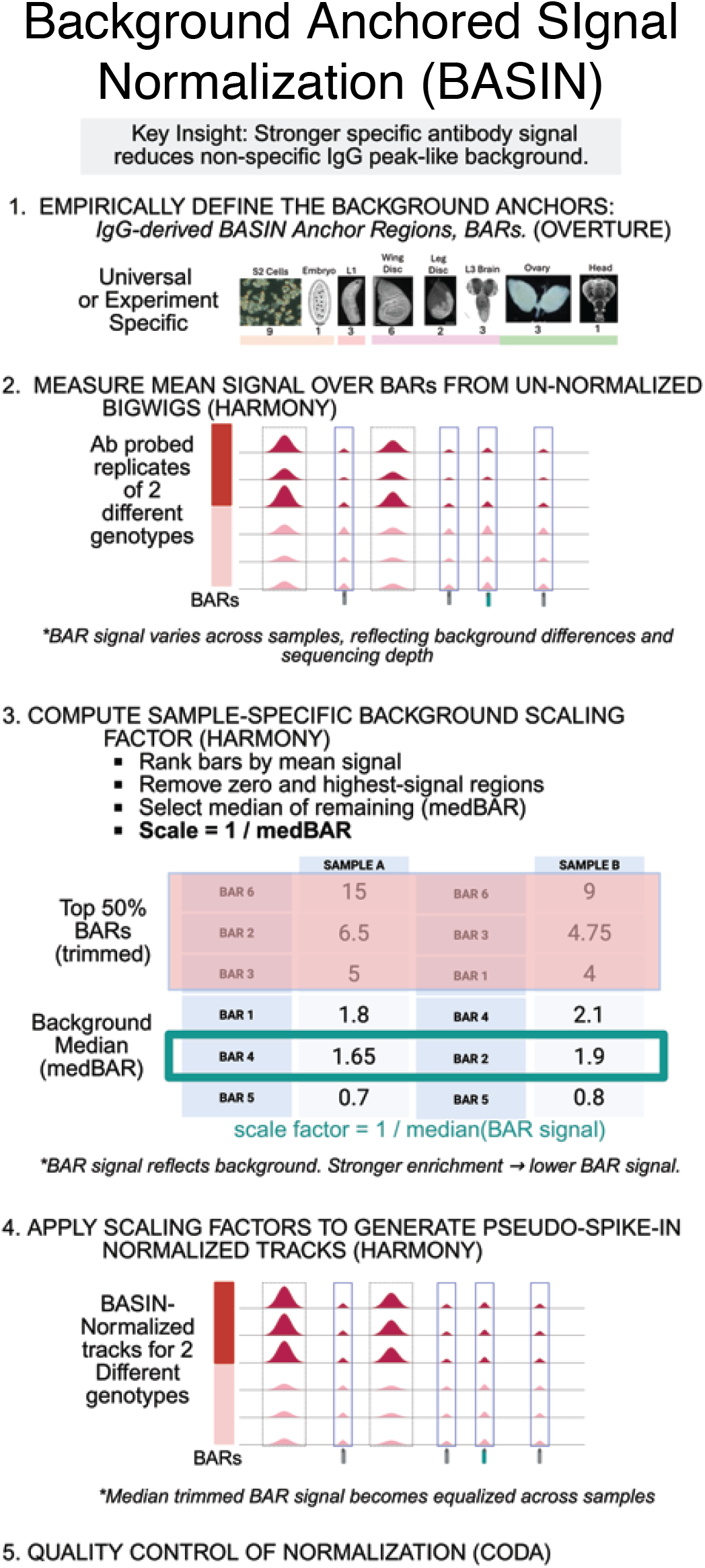
BASIN and the details of the HARMONY normalization workflow. BASIN is comprised by three pipelines — OVERTURE (Fig. 3), HARMONY, and CODA (Fig. 7) — for anchor region discovery, sample normalization, and post-normalization quality control, respectively. This figure details the HARMONY workflow. **(1)** HARMONY normalizes each sample using its BAM file and a BAR (BASIN Anchor Region) panel generated by OVERTURE, using either a universal or an experiment-specific panel. Images show the tissues and cell types that contributed IgG datasets to the universal *Drosophila melanogaster* panel, with the number of datasets beneath each; colored bars group them by developmental stage. **(2)** Mean coverage at each BAR is summarized from raw, library-size-uncorrected bigWig files. Schematic tracks for three replicates of each of two genotypes show that raw BAR signal varies across samples, reflecting both genuine differences in background and differences in sequencing depth. Blue boxes mark BARs; dashed boxes mark domains of specific signal that are not BARs. **(3)** BAR values are ranked in descending order; the top fraction (the drop threshold) is trimmed along with BARs showing zero signal across all samples, and the median of the remaining values (medBAR) is calculated. The scale factor for that sample is 1/medBAR. A simplified example using hypothetical BAR values from two samples illustrates the ranking, trimming (here, the top 50%), and median steps. **(4)** HARMONY regenerates a bigWig directly from each sample’s BAM using bamCoverage, applying that sample’s scale factor at the coverage-generation step (–– scaleFactor). Schematic tracks show BASIN-normalized signal for two genotypes at the locus shown in (2); the median signal over retained BARs is equal across samples by construction. **(5)** Output files are passed to CODA for quality control.

The trimming step employed in HARMONY is conceptually similar to the trimmed mean of M-values (TMM) method employed for RNA-seq normalization or reads-in-peak methods (Robinson & Oshlack, 2010; Ross-Innes et al., 2012; Stark & Brown, 2011). Many other endogenous normalization approaches define or filter reference regions using prior genomic or biological annotation, including blacklist/mappability filters, invariant CTCF sites, promoters, or other presumed stable functional elements (Cao et al., 2026; de Mello et al., 2024; Oh et al., 2026; Xiang et al., 2020). In BASIN, trimming is instead applied empirically without reference to gene annotation, so anchors contaminated by true signal are removed on the basis of their amplitude rather than their genomic identity. We reasoned that this approach would be less likely to bias normalization behavior for features residing in euchromatic versus heterochromatic compartments. Earlier ChIP-seq background-based normalization approaches such as NCIS similarly recognized that normalization-factor estimation requires balancing contamination by true enrichment against the variance incurred when too little background signal is retained (Liang & Keleş, 2012). BASIN addresses the same general tradeoff through empirical BAR– panel construction and robust upper-tail trimming, but deliberately uses recurrent high-count IgG-associated background rather than low-count genomic bins. The BASIN scale factor is applied directly to the original BAM file when normalized bigWigs are generated using bamCoverage with the ––scaleFactor argument, rather than by rescaling an existing coverage track. The resulting file expresses signal relative to that sample’s own empirical background.

A single BASIN scaling operation performs two functions simultaneously since both raw coverage at a genomic position and the sample’s median BAR coverage (medBAR) scale with sequencing depth (Fig. S1). Dividing raw coverage by medBAR therefore cancels their common depth factor, expressing signal as a ratio to the sample’s background anchor. Under the assumption that BARs represent a genotype-independent background, this ratio provides a common scale for between-sample comparisons. Sequencing-depth adjustment and background anchoring are thus achieved through a single operation, without a separate library-size normalization step. Because the BASIN scale factor is defined such that medBAR equals unity after scaling, convergence of anchor medians is an algebraic consequence of normalization rather than independent evidence of its validity. We therefore use post-normalization anchor convergence as a pipeline integrity check, not as a test of whether BARs provide a biologically stable reference.

### OVERTURE identifies reproducible anchor regions from public and experiment-specific IgG datasets

The utility of an endogenous normalization reference is only as good as the genomic regions are chosen for reference. We therefore developed OVERTURE, a discovery and region evaluation pipeline that identifies genomic regions recurrently recovered in IgG controls and then benchmarks candidate panels for the properties required of a background anchor reference (Fig. 3A). OVERTURE can be used in two modes. In a universal mode, candidate panels are derived from a heterogeneous collection of public and in-house IgG datasets, with the goal of identifying regions that behave as reproducible background across many biological and technical contexts. In an experiment-specific mode, candidate panels are derived from IgG controls generated within a single experimental system, providing a context-matched alternative. A validated universal BAR panel for *Drosophila melanogaster* is distributed with the software package.

**Figure 3.**
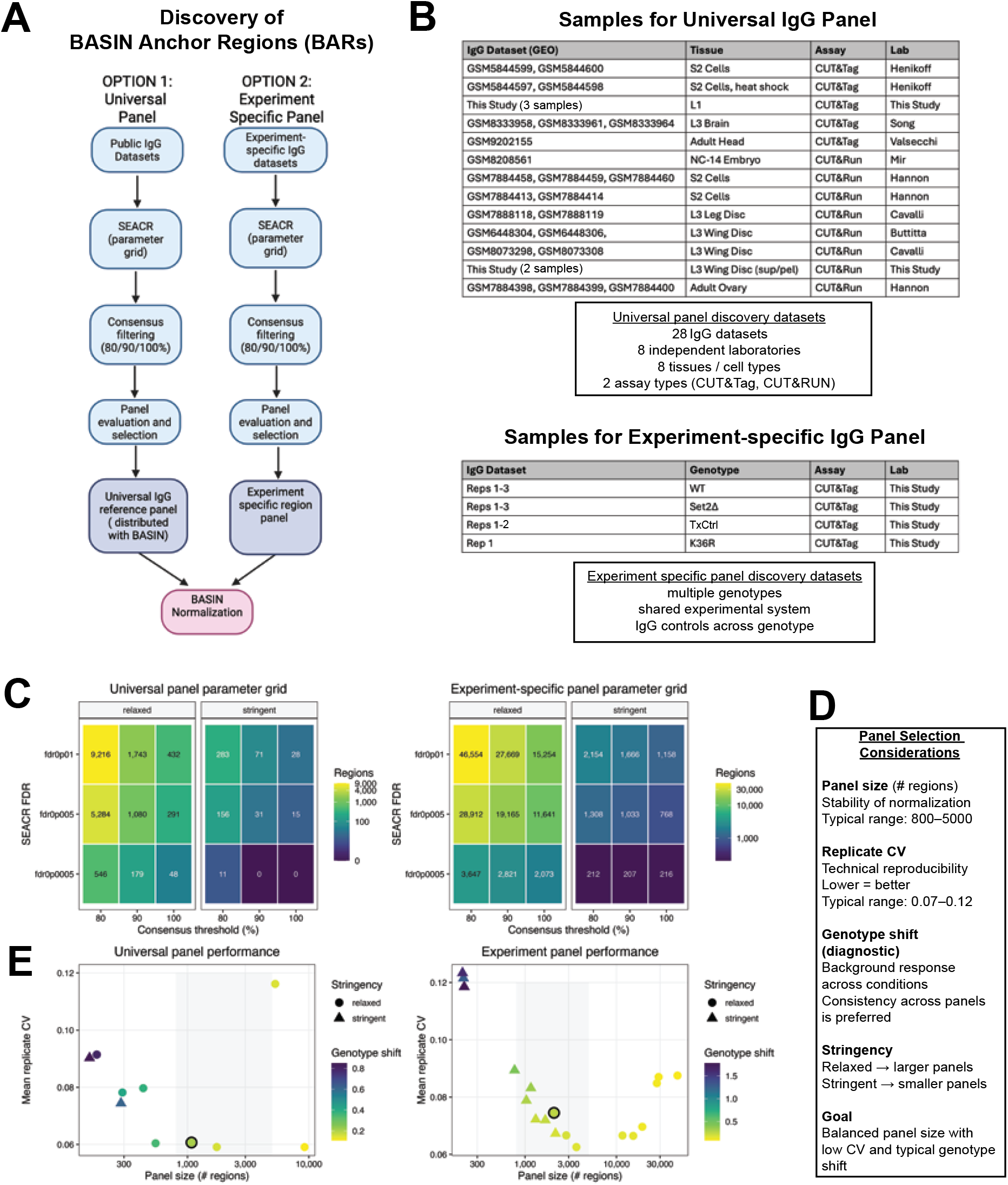
OVERTURE identifies reproducible BASIN Anchor Regions (BARs) from public and experiment-specific IgG datasets. **(A)** Overview of the OVERTURE BAR discovery workflow. Public IgG CUT&Tag/CUT&RUN or experiment-specific IgG control datasets are used to identify genomic regions that consistently accumulate IgG signal. Universal anchor panels are generated from diverse public and in-house datasets, whereas experiment-specific panels are generated from IgG controls associated with a single experimental system. Candidate regions are identified by parameter sweeps over SEACR threshold, SEACR stringency, and consensus threshold (recurrence in 80%, 90%, or 100% of datasets), evaluated for normalization performance, and compiled into anchor panels used by HARMONY during BASIN normalization. **(B)** Datasets used for anchor panel discovery in *Drosophila melanogaster*. Universal panel discovery incorporated 28 publicly available and in-house generated IgG datasets spanning eight independent laboratories, eight tissue or cell types, and two assay types (CUT&Tag and CUT&RUN). Experiment-specific panel discovery used nine IgG controls generated within the same experimental system across four genotypes (WT, Set2Δ, TxCtrl, and K36R) to identify anchors represented in the specific experiment elaborated in Figure 4. **(C)** Number of anchor regions retained across a parameter sweep of SEACR threshold (0.01, 0.005, or 0.0005), SEACR stringency (relaxed or stringent), and consensus threshold (80%, 90%, or 100%), for universal (left) and experiment-specific (right) panel construction, after exclusion of regions shorter than 50 bp; heatmap color corresponds to the number of regions retained under each condition. **(D)** Criteria used for final panel selection: region count, technical reproducibility (replicate CV), genotype shift, and stringency. Larger panels generally provide increased stability, while more stringent filtering yields smaller, higher-confidence anchor sets. Genotype shift is diagnostic, but choosing a panel with genotype shift similar to other panels is desirable. The selected parameter combination balances panel size with low replicate CV and a genotype shift typical of other panels to generate the BAR files used by HARMONY. Shading indicates the recommended operating range (800–5,000 BARs; mean replicate CV <0.12). **(E)** Relationship between anchor panel size (filtered for panels with >50 regions) and mean replicate CV for universal (left) and experiment-specific (right) panel discovery, evaluated on the six-target L1 CUT&Tag datasets for H3K36me2, H3K36me3, panH4ac, H3K27ac, RNAPII-S5P, and RNAPII-S2P. The shaded band marks the 800–5,000-region range used for panel selection. Each point represents one parameter combination; marker shape denotes SEACR stringency, and color indicates medBAR genotype shift. The outlined point is the selected panel used in subsequent figures.

For universal panel discovery in *D. melanogaster*, we compiled 28 IgG datasets spanning eight independent laboratories, eight tissues or cell types, 4 developmental stages, and both CUT&Tag and CUT&RUN assays (Fig. 3B). By contrast, experiment-specific panel discovery used IgG controls collected across genotypes within the first larval stage (L1) H3K36 experimental system analyzed below, providing a within-experiment discovery strategy for comparison. These two modes are conceptually complementary: universal discovery prioritizes robustness across contexts, whereas experiment-specific discovery prioritizes relevance within a shared experimental background. However, experiment-specific panels may be more susceptible to inclusion of regions of experimental system-specific coverage, such as accessible regions.

Candidate panels were generated by calling peaks in individual IgG datasets and then constructing consensus regions across samples using parameter sweeps over SEACR (Meers et al., 2019a) stringency and consensus thresholds requiring recurrence in 80%, 90%, or 100% of input datasets. OVERTURE also produces versions of these consensus files where regions less than 50bp are filtered out, to ensure that potential BARs have a sensible lower bound on size. This produced a broad range of candidate panels whose region counts varied substantially across the parameter grid (Fig. 3C). As expected, relaxed peak-calling settings and lower consensus thresholds yielded the largest panels, whereas stringent settings and higher consensus requirements produced progressively smaller panels and, in some cases, few or no candidate regions. At matched parameter settings, experiment-specific discovery generally produced larger candidate panels than universal discovery, consistent with the greater recurrence expected within a relatively homogeneous set of related datasets.

OVERTURE next evaluates candidate panels using user-provided validation datasets and three complementary diagnostics: panel size, replicate reproducibility, and genotype-dependent background behavior (Fig. 3D). For the universal dm6 analysis, we used a six-target L1 CUT&Tag evaluation dataset comprising H3K36me2, H3K36me3, H3K27ac, and pan-H4ac histone modifications, as well as RNAPII-S2P and RNAPII-S5P phosphorylations, each measured across biological replicates and genotypes. After panel selection, this universal panel was further used and assessed on other datasets: H3K9me3, H3K27me3, and pan-H4ac wing disc CUT&RUN, and H3K27me3 and H3K4me1 CUT&Tag (detailed below). Panel size determines whether enough anchors are available to generate a stable trimmed-median estimate of background signal. Replicate coefficient of variation (CV) quantifies technical reproducibility of the anchor-derived background estimate within genotypes or conditions. OVERTURE also reports medBAR genotype shift, defined as the change in trimmed median BAR signal between a test genotype and the reference genotype. Importantly, genotype-dependent changes in medBAR are not inherently evidence of panel failure. Because sequencing data are compositional, a genuine global change in specific signal can alter the relative representation of nonspecific background, producing coordinated background shifts across many candidate panels. medBAR genotype shift is therefore interpreted as a diagnostic of whether a candidate BAR panel exhibits the expected background titration behavior, not as a quantity that should simply be minimized toward zero (See OVERTURE documentation).

Plotting replicate CV against panel size across the parameter grid reveals the expected tradeoff between panel breadth and technical stability (Fig. 3E, FigS2A). Very small panels (<500 BARs) were less likely to pass, and are more susceptible to fluctuations at individual BARS, whereas very large panels may incorporate more regions that are susceptible to condition related variability (Fig. 3E, FigS2A). Panels containing at least 200 BARs were evaluated; based on these results, we recommend a working range of 800–5,000 BARs and a mean replicate CV below 0.12 (shaded in Fig. 3E). Restricting the analysis to candidate panels containing more than 50 BARs shifted the distribution toward smaller median BAR lengths (Fig. S2B). These candidate panels were then evaluated for medBAR genotype shifts across genotypes and test dataset marks (FigS2C). Shared shifts observed across many candidate panels are most consistent with dataset-level background titration due to global shifts in signal, whereas unusually panel-specific or discordant shifts may indicate contamination by genotype-responsive specific signal. To make this distinction more explicit, we additionally summarized “genotype-shift typicality” across passing universal panels (FigS2D,E). This metric assesses whether the medBAR genotype shifts for different test genotypes and targets tend to remain close to the panel median as a whole, or whether behavior diverges significantly from other panels (FigS2D). Importantly, genotype-shift typicality does not penalize panels where BARs scale globally in response to global changes in target signal. In this analysis, the selected universal panel, defined using relaxed SEACR peak calling (FDR 0.005), 90% consensus, and exclusion of regions shorter than 50 bp, behaved representatively relative to the ensemble of passing panels, ranking first by mean typicality distance and third by median typicality distance (Fig. S2E). Thus, the selected panel was not chosen because it artificially minimized genotype shifts, but because it combined acceptable size and reproducibility with genotype-shift behavior that was broadly consistent with other well-performing candidates.

Together, these analyses show that OVERTURE can identify BAR panels that are both technically stable and biologically interpretable. Universal discovery prioritizes regions that recur across diverse IgG controls and therefore are likely to generalize across experiments, whereas experiment-specific discovery provides a context-matched alternative when appropriate controls are available. The resulting workflow supplies an explicit and quantitative framework for selecting endogenous anchor regions rather than treating background normalization as an implicit assumption.

### BASIN recovers the expected depletion of H3K36me3 in Set2Δ and H3K36R mutants

A normalization method that preserves global differences should be tested against a perturbation whose global effect is known independently. Loss of Set2, the principal H3K36 trimethyltransferase in *Drosophila*, and substitution of H3K36 to arginine (H3K36R) both reduce H3K36me3 genome-wide (Larschan et al., 2007; Meers et al., 2017; Sultana et al., 2026). To confirm the degree of H3K36me3 in our developmentally earlier larval stage 1 (L1) experimental timepoint, we performed Western blotting on mutant and control lysates at this stage. This confirmed global reduction of bulk H3K36me3 in our larval 1 stage (L1) experimental system both when normalized to an H4 loading control (Fig. 4A), and total protein (Fig.S3). The two perturbations differ in severity: Set2Δ removes trimethylation (∼60% of control by western; Fig. 4A) while leaving the residue intact, whereas H3K36R abolishes all modification states of H3K36, leaving essentially no specific signal for the antibody to recover (Fig. 4A). The reduction of bulk H3K36me3 at this stage is subtler than that reported for older animals, possibly due to residual maternal contribution or activity of another H3K36 methyltransferase at L1. In contrast, K36R represents the most extreme global shift under which a normalization method can be tested, and its behavior throughout this section reflects that. We profiled H3K36me3 by CUT&Tag in Set2Δ and H3K36R L1 larvae alongside their matched controls; the alleles and genetic systems used to generate these mutants have been described previously (Larschan et al., 2007; McKay et al., 2015; Salzler et al., 2024). Because these libraries carried a high duplication rate (∼86% on average;, Table T1, highlighted in yellow), we performed our primary analyses on unique fragments.

**Figure 4.**
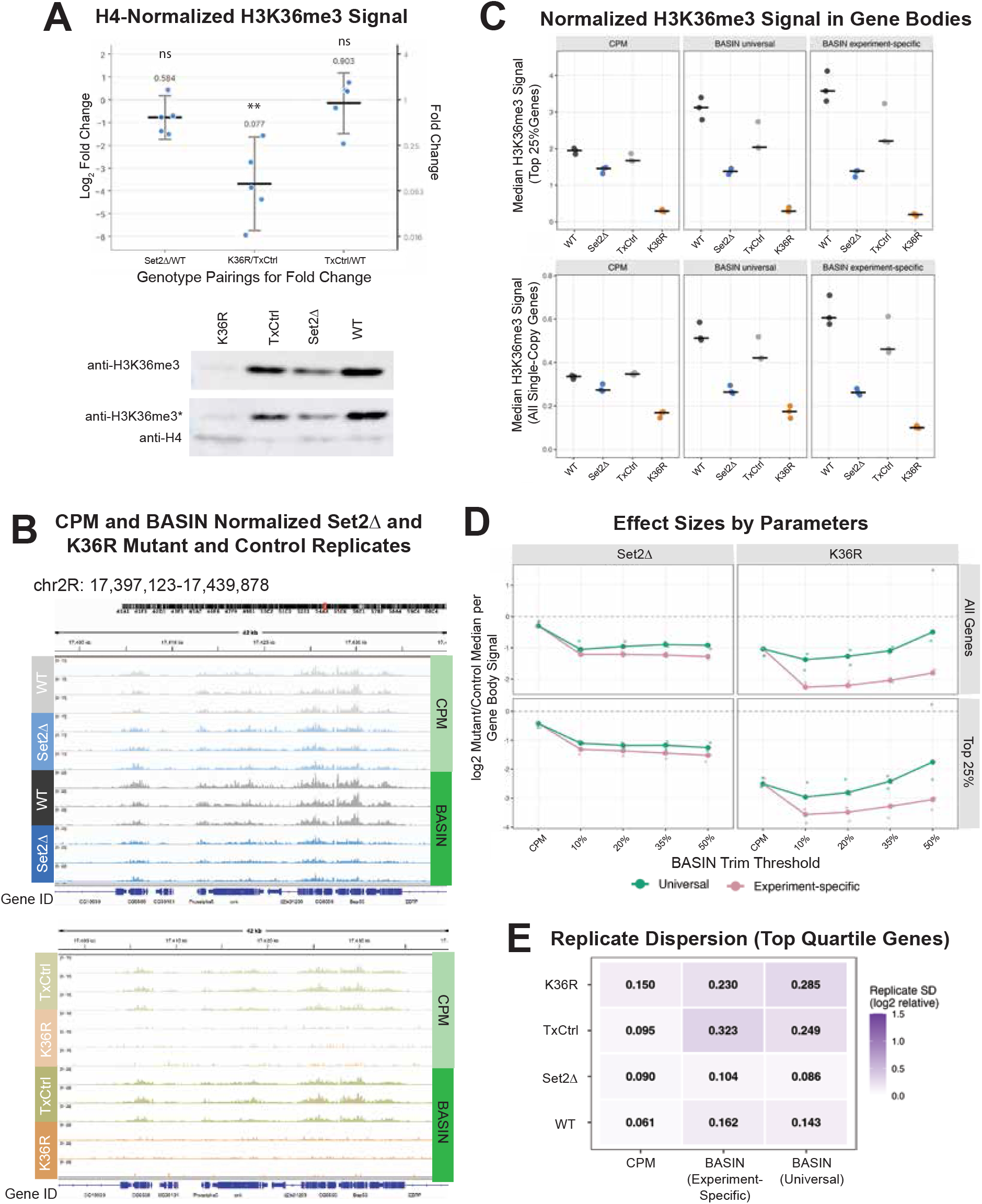
BASIN normalization recovers expected H3K36me3 depletion in Set2Δ and H3K36R mutants after duplicate removal. **(A)** Representative H3K36me3 western blot validating global reduction of H3K36me3 abundance in Set2Δ and K36R mutants. H3K36me3 was compared to an H4 loading control for 5 biological replicates per genotype. Anti-H3K36me3* indicates residual H3K36me3 signal left on blots when re-probed for H4 because blots were not stripped (see methods). For quantification, H3K36me3 band intensities were normalized to pan-H4 loading-control intensities within each lane. Normalized intensities were expressed as both fold changes and log2 fold changes relative to the corresponding matched control for each biological replicate. The geometric mean (GM) of each comparison is calculated (mutant/control) or (control/control) and GM and statistical significance is reported above each column. Error bars indicate the 95% confidence interval. Statistical significance was calculated using two-tailed paired t-tests on log2 fold-change values (*p < 0.05; **p < 0.01; ns, not significant). **(B)** Representative genome-browser views (chr2R:17,397,123–17,439,878) of deduplicated H3K36me3 profiles for Set2Δ and H3K36R mutant replicates and matched controls (WT and TxCtrl, respectively; three replicates per genotype), following CPM or BASIN (drop0.20) normalization. Tracks are displayed on identical scales within each normalization method. **(C)** Median normalized H3K36me3 signal across gene bodies of the top quartile of H3K36me3-marked genes and all single-copy genes after duplicate removal, under CPM, BASIN (drop0.20) with the universal panel, or BASIN (drop0.20) with the experiment-specific panel. Points represent individual biological replicates; horizontal bars denote the median across replicates. Normalization units are not comparable between normalization methods. **(D)** Relative effect sizes for Set2Δ (left) and K36R (right) across BASIN anchor-drop thresholds after duplicate removal, for all genes (top) and the top quartile of H3K36me3-marked genes (bottom). Values are the log₂ ratio of mutant gene-body signal to the median of the matched control; small points are individual replicates and large points the median across replicates. CPM is plotted at the left of each panel, and the dashed line marks no change. Estimates from the universal (green) and experiment-specific (pink) panels are stable and concordant across thresholds for Set2Δ and diverge for K36R. **(E)** Replicate dispersion for the top quartile of H3K36me3-marked genes following duplicate removal, for CPM and for BASIN (drop0.20) with the experiment-specific or universal panel. Replicate dispersion was calculated as the standard deviation across biological replicates of log2-transformed sample signal relative to the median of the matched control replicates for each normalization. Sample signal was defined as the median normalized H3K36me3 signal across this gene set. The full threshold series and the all-genes analysis are shown in Supplementary Figure S4A.

Genome-browser views illustrate the practical consequence of read-depth normalization on conditions generating a global decrease (Fig. 4B). Under CPM normalization, WT and Set2Δ tracks are nearly indistinguishable across gene-rich intervals. The same samples normalized by BASIN separate clearly, with Set2Δ reduced relative to WT across the interval. The K36R reduction is apparent under both normalizations, as expected given the near-complete loss of signal in that genotype. We also tested our experiment-specific panel generated using IgG controls from all experimental and control genotypes at L1 stage (Fig. 3E,4C-E) side by side for a quantitative comparison. Because our transgenic histone replacement system introduced known histone array copy-number differences system 100 copies for WT and Set2Δ; ∼12 copies for TxCtrl and H3K36R) (McKay et al., 2015), we manually removed intervals overlapping with the histone array from the universal panel bringing the final number of BARs from 1080 to 856. Of note, the experiment-specific panel only required removal of 3 intervals. Quantification of median signal across the gene bodies of the most highly H3K36me3-marked quartile of genes demonstrates large reductions between mutant and control genotypes for both BASIN panels with negligible difference between WT and Set2Δ detectable by CPM (Fig. 4C). Because the two normalizations express signal in different units, absolute values are not comparable between methods, thus only the relative separation between genotypes is informative.

Effect-size estimates were stable across anchor-drop thresholds for Set2Δ, and universal and experiment-specific anchor panels produced concordant estimates across a broad range of thresholds (Fig. 4D). This is a useful result for practical application of BASIN, since it indicates that users need not tune the drop threshold precisely. Reassuringly, the effect size in the Set2Δ mutant was commensurate to the mean value observed by western blot (Fig.4A,D). K36R behaved differently in two respects. The two anchor panels diverged substantially, with the experiment-specific panel producing consistently larger estimated depletion and less threshold dependence than the universal (Fig4D). Where no specific signal remains, normalization behavior is determined almost entirely by anchor composition, and two properties of the panels become consequential. The universal panel comprises fewer regions than the experiment-specific panel (Fig. 3C), so aggressive trimming leaves proportionally fewer values supporting the median and medBAR becomes correspondingly less stable. Panel composition may also contribute: OVERTURE’s consensus filter selects regions on the basis of reproducible peak presence rather than signal magnitude, so an experiment-specific panel derived within a single tissue and developmental stage can retain regions whose background behavior is context-dependent, whereas the universal panel is filtered for recurrence across tissues and assay chemistries. With respect to an experiment-specific panel, a genotype or condition with globally altered chromatin accessibility would shift anchor signal uniformly, and upper-tail trimming cannot correct for this. Trimming operates on rank within a sample and is insensitive to a uniform multiplicative shift, even when effective against subsets of anchors containing real signal. This vulnerability is shared by other endogenous-reference methods and is not equally shared by all normalization strategies (see Discussion).

Replicate dispersion was quantified, within each genotype group, as the standard deviation across biological replicates of log2 H3K36me3 signal relative to the median of the matched control replicates for each normalization method. Using this metric, dispersion over the top quartile of genes was generally lower under CPM than under BASIN for most genotypes (Fig. 4E), as expected because CPM constrains all libraries to the same total signal and therefore suppresses variation in global signal abundance. Across moderate trim thresholds, the universal and experiment-specific anchor panels yielded broadly similar replicate dispersions across genotypes (Fig. 4E; Fig. S4). In K36R, however, performance of the universal panel deteriorated markedly at drop0.50 (Fig. S4), the same trim level at which effect-size estimates drifted toward zero (Fig. 4D). One likely explanation is that in K36R, where true H3K36me3 signal is largely absent, aggressive trimming removes a disproportionate fraction of the BAR values still providing stable support for medBAR. This effect is expected to be most severe for the smaller universal panel, leaving the background estimate to be determined from a reduced and noisier residual set. Consistent with this interpretation, deduplication substantially reduced universal-panel dispersion in K36R at working thresholds while leaving the experiment-specific panel comparatively unchanged (Fig. S4). Because duplication rates were similar across genotype groups, this disproportionate improvement is consistent with duplicated background reads contributing more strongly once anchor support becomes sparse. We therefore recommend conservative trimming, with a default drop threshold of 0.10–0.20, and caution that trimming the universal panel beyond 0.35 is inadvisable in samples approaching complete loss of the profiled target.

For robustness, we repeated these analyses with duplicate reads retained (Fig. S5A–D), which additionally allows comparison with spike-in normalization. Spike-in reads for these experiments derive from E. coli carryover in the pA-Tn5 preparation (Meers et al., 2019b); carryover material is introduced as a fixed amount per reaction and yielded marginal read counts (Table T1, highlighted in blue). This yielded too few reads after deduplication to estimate a scale factor, so this comparison is restricted to the duplicate-retaining analysis. The relative behavior of CPM and BASIN was consistent between the two modes of duplicate handling (Fig 4A-E; S5A,B,D). Spike-in recovered the expected global changes, but showed greater replicate variability than BASIN across nearly all conditions examined. At a conservative drop threshold of 0.10, the experiment-specific panel gave lower replicate dispersion than spike-in in every genotype and both gene sets, and the universal panel did so in all but one. Differences were often large: for all genes at drop0.20, replicate dispersion for the universal panel versus spike-in was roughly four-fold (Fig S4). The one condition in which spike-in was more stable than BASIN was the universal panel in K36R, where anchor-median instability is already established. Because the same samples underlie every normalization compared, these differences are attributable to the reference estimate itself. Spike-in therefore performs relatively best in precisely the genotype where the universal panel is least stable, and offers no advantage elsewhere. Given that its reference in this experiment derives from carryover rather than a controlled spike-in protocol, we treat spike-in as a supplementary comparison rather than as a benchmark against which BASIN is evaluated.

### BASIN preserves input-independent signal across cell-number titrations

The H3K36me3 experiment tests whether BASIN can detect a genuine global change when one is present. The complementary question is whether BASIN reports no change when no biological change should exist. A cell-input titration series, in which chromatin state is held constant while the amount of input material varies, provides this control. It also offers the advantage that the expected behavior of each normalization method can be specified quantitatively in advance.

Under these conditions, CPM and BASIN are both expected to return input-independent normalized signal because their scaling references are derived internally from the assayed sample. In log2-log2 space, this corresponds to an expected slope of zero. Spike-in normalization, by contrast, derives its reference externally. Because a fixed amount of reference chromatin is added to each reaction while sample input varies, the recovered reference should ideally vary inversely with sample input, such that spike-in-normalized signal scales in direct proportion to the amount of sample, corresponding to an expected slope of one (Fig. 5A). These are predictions about normalization behavior, not about the underlying biology, and each method can therefore be evaluated against its own theoretical expectation.

**Figure 5.**
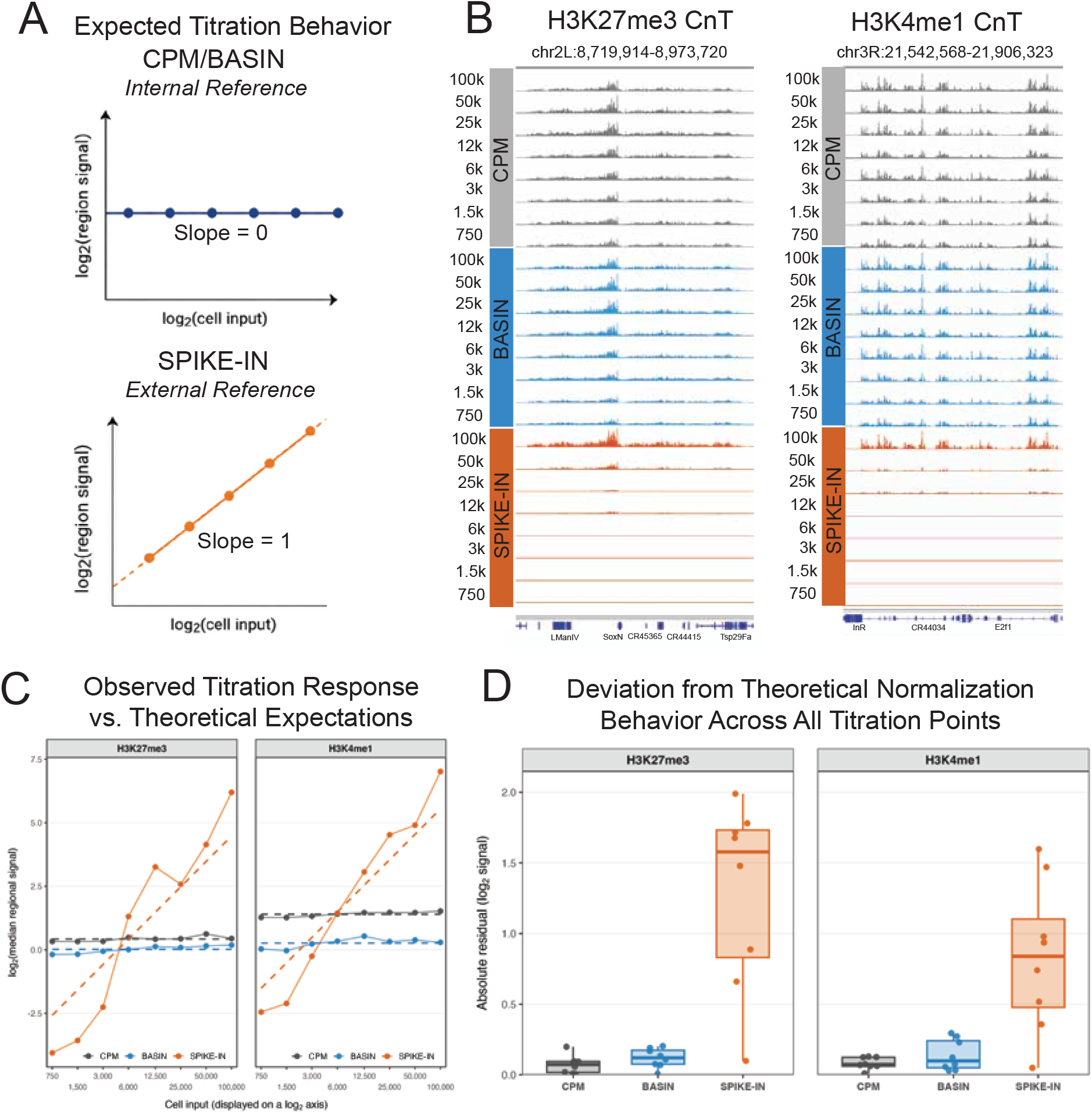
BASIN preserves input-independent signal across a cell-input titration, whereas spike-in normalization deviates from its own expected response. (A) Theoretical normalization responses expected in a titration experiment in which cell input changes but the underlying chromatin state remains constant. CPM and BASIN use internal genomic signal as the normalization reference and are therefore expected to maintain constant normalized regional signal across cell inputs, corresponding to a slope of 0 in log₂–log₂ space. Spike-in normalization uses an external reference and is expected to produce normalized signal proportional to cell input, corresponding to a slope of 1. (B) Representative genome-browser views of H3K27me3 (left; chr2L:8,719,914–8,973,720) and H3K4me1 (right; chr3R:21,542,568–21,906,323) CUT&Tag titration series spanning eight inputs from 100,000 to 750 *Drosophila* S2 cells. Each reaction also received 200 human K562 cells as an exogenous reference. Tracks are shown for CPM normalization, BASIN normalization at the 0.20 anchor-drop setting, and spike-in normalization. A fixed display scale was used within each normalization-method block. CPM and BASIN preserve broadly stable signal profiles across the titration series, whereas spike-in-normalized signal declines sharply at lower cell inputs, as expected from panel A. **(C)** Observed median regional signal compared with the theoretical response for each normalization method. Median signal was calculated across the top 25% of regions for each histone modification. Cell input and regional signal are displayed on log₂ scales. Solid lines and points show observed values; dashed lines show theoretical models with slopes fixed at 0 for CPM and BASIN and 1 for spike-in normalization. For each histone modification and normalization method, the intercept was estimated across all eight titration points. Signal values between each method are not comparable, only slopes should be compared. **(D)** Absolute deviation of each titration point from the corresponding fixed-slope theoretical model. Each point represents one cell-input level; boxplots summarize the eight absolute residuals for each normalization method. Lower values indicate closer agreement with expected normalization behavior. CPM and BASIN closely follow the expected input-independent response for both histone modifications, whereas spike-in normalization shows substantially larger deviations.

We analyzed H3K4me1 and H3K27me3 CUT&Tag data across an eight-point titration spanning 100,000 to 750 *Drosophila* S2 cells, with 200 K562 human cells added to each reaction as an exogenous reference. Browser views showed that CPM– and BASIN-normalized tracks remained broadly stable across the series, whereas spike-in-normalized signal declined at lower inputs, consistent with the qualitative behavior expected for a method that preserves input– dependent changes in signal (Fig. 5B). To assess these trends quantitatively, we measured median signal over the most enriched quartile of regions for each modification and plotted these values against cell input together with the corresponding fixed-slope theoretical models (Fig. 5C). CPM and BASIN tracked their expected input-independent behavior closely across the full titration range, whereas spike-in deviated substantially from its own expected linear response. We quantified this agreement by calculating residuals (deviation from expected at each point) from each method’s theoretical model and summarizing their absolute values (Fig. 5D). For both H3K4me1 and H3K27me3, CPM and BASIN showed substantially smaller deviations from expectation than spike-in. Thus, the limitation in these experiments was not simply that spike-in behaved differently from the internal-reference methods, which is expected, but that it failed to behave as spike-in normalization itself predicts it should.

Read composition across the titration series helps explain this discrepancy (Table T1). Because the human genome is approximately 30-fold larger than the *Drosophila* genome, the 200 K562 cells added to each reaction contribute DNA equivalent to roughly 6,000 fly cells. In principle, this should provide ample exogenous reference material across the series, from approximately 6% of total DNA at 100,000 fly cells to more than the sample itself at the lowest inputs. However, the processed human reference counts were not proportional across the titration range. Although total library depths were similar, fly read counts varied only approximately threefold across the 133-fold range of cell input, whereas human reference reads varied 332-fold for H3K4me1 and more than 1,100-fold for H3K27me3. At 750 cells, reference reads constituted 8.1% and 5.0% of the respective libraries, but at 100,000 cells they accounted for only 0.013% and 0.004%, corresponding to just 975 and 178 reads. These values are far below those predicted from the input proportions, and the shortfall is concentrated at the high-input end of the series. If the processed reference counts scaled ideally, the product of reference reads and fly cell input would remain constant across the titration. Instead, this quantity varied by approximately fivefold for H3K4me1 and sixteenfold for H3K27me3, with the lowest values at the highest fly-cell inputs. Accordingly, the processed human reference declined more steeply than the inverse of input, causing spike-in-normalized signal to rise more steeply than the theoretical slope of one.

There are several non-exclusive explanations for the discordance of spike-in normalization from it’s theoretical prediction. First, simple pipetting of cells in suspension may have been non-uniform. Second, the exogenous reference material may not have been recovered proportionally across the titration series, particularly at higher fly-cell inputs. Third, upstream processing effects—including mapping, species assignment, or filtering—may have distorted the processed human read counts used for spike-in normalization. Any or all mechanisms could cause the processed spike-in counts to deviate from the theoretical spike-in response. Importantly, these contingencies are intrinsic to the practical performance of spike-in normalization, which depends on accurate recovery and quantification of an exogenous reference population. By contrast, CPM and BASIN do not rely on cross-species reference recovery or species-assignment workflows and are therefore not subject to these particular sources of error. In fact, the very same titration series normalized with BASIN behaved almost perfectly according to theoretical expectations. Taken together, these results indicate that in this titration series CPM and BASIN more faithfully matched their expected behavior than spike-in normalization did.

### BASIN recovers H3K9me3 signal structure where anchor and target regions overlap

The preceding experiments test BASIN under conditions favorable to it, in that the anchor regions and the profiled modification occupy largely distinct parts of the genome. The assumption that anchors represent background independent of the target signal is most likely to fail when the two overlap. We therefore sought a dataset in which that overlap is unavoidable.

H3K9me3 profiling in L3 wing disc provides several such difficulties simultaneously (Fig. 6A): the profiled modification occupies heterochromatin, which is also where anchor regions are concentrated (Fig. 6B); the tissue is one in which exogenous spike-in normalization is technically difficult due to the inability to obtain cell counts without dissociation; the genotype groups are sex-skewed; and H3K9me3-enriched regions overlap substantially with repetitive sequence that is difficult to map uniquely.

**Figure 6.**
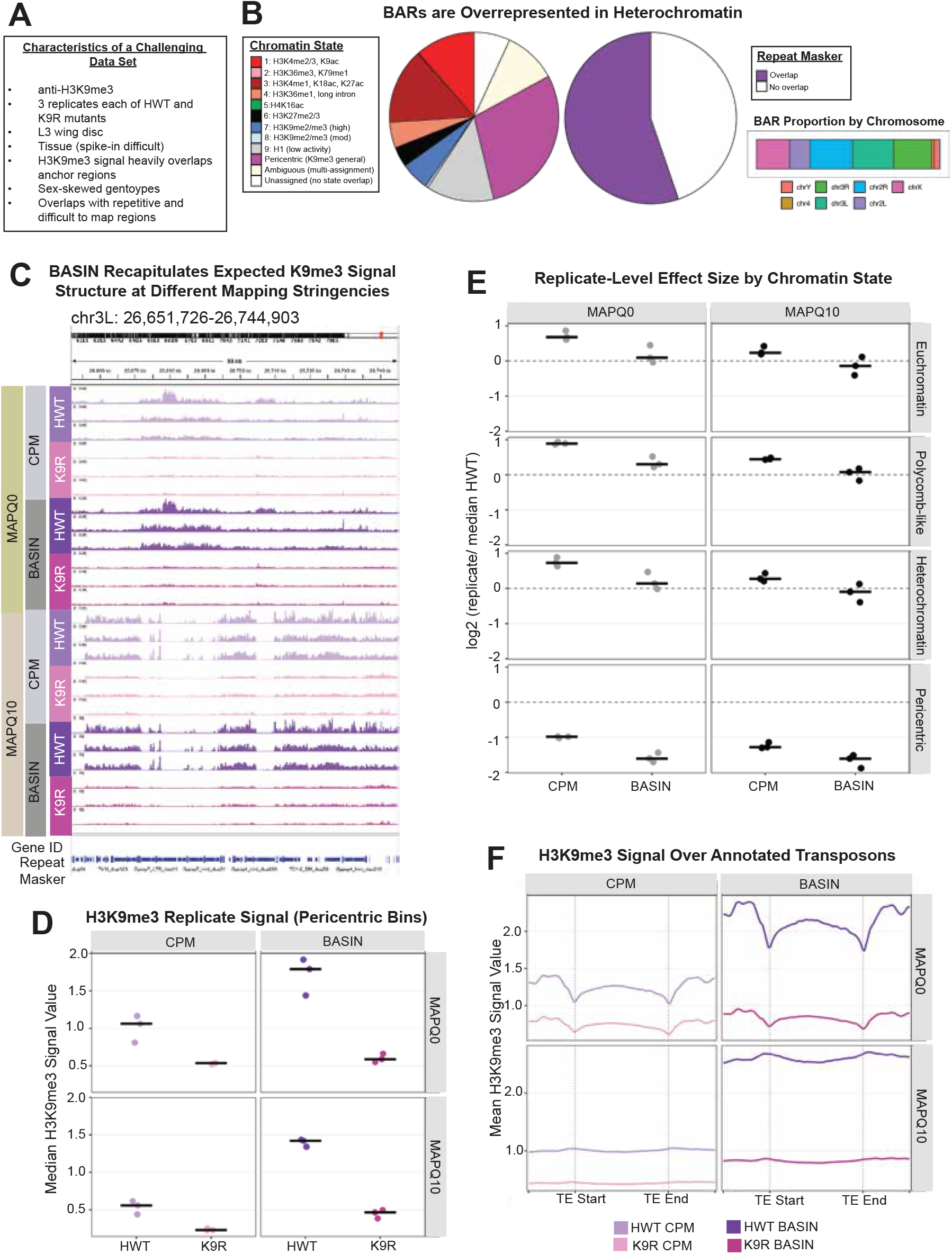
BASIN recovers expected H3K9me3 signal structure in a challenging heterochromatic dataset. **(A)** Summary of experimental features that make this dataset a stringent test of BASIN normalization: an antibody (H3K9me3) whose target regions substantially overlap the anchor regions used for normalization; profiling in L3 wing disc, a tissue in which exogenous spike-in normalization is technically difficult; sex-skewed genotype composition (histone-wild-type HWT and K9R histone-replacement genotypes, three replicates each); and substantial overlap between H3K9me3-enriched regions and repetitive, difficult-to-map sequence. **(B)** BASIN Anchor Regions (BARs) are overrepresented in heterochromatin. Left pie: distribution of chromatin states (Kharchenko 2011, “pericentric” inferred by coordinates) overlapping BARs, showing enrichment for H3K9me2/3-associated and pericentric states relative to euchromatic states. Right pie: fraction of BARs overlapping RepeatMasker-annotated repetitive elements. Right bar: proportion of BARs by chromosome, colored by chromosome identity. **(C)** Representative genome-browser view of H3K9me3 signal for HWT and K9R replicates at a pericentromeric heterochromatin locus (chr3L:26,651,726–26,744,903), shown at two mapping-quality thresholds (MAPQ0, retaining multimapping reads consistent with BAR panel discovery, and MAPQ10) for CPM and BASIN (drop0.20) normalization. BASIN-normalized tracks recapitulate the expected loss of H3K9me3 signal in K9R relative to HWT at both mapping stringencies. Track scales are auto-scaled separately for each method/MAPQ combination. **(D)** Median H3K9me3 signal in pericentric bins for HWT and K9R replicates following CPM or BASIN normalization, at MAPQ0 (top) and MAPQ10 (bottom). Each point represents one biological replicate; horizontal bars indicate the median across replicates. **(E)** Replicate-level effect sizes, expressed as log₂(replicate signal / experiment-group median HWT signal), for CPM– and BASIN-normalized data, stratified by chromatin state (euchromatin, Polycomb-like, heterochromatin, and pericentric). Columns show MAPQ0 (left) and MAPQ10 (right). Points are individual K9R replicates, horizontal bars indicate the median, and the dashed line marks no change. **(F)** Mean H3K9me3 signal over annotated transposable elements, scaled to a common length and flanked by their start and end coordinates, for HWT and K9R replicates following CPM or BASIN normalization at MAPQ0 (top) and MAPQ10 (bottom). BASIN-normalized signal shows the expected relative enrichment of H3K9me3 in HWT versus K9R over transposable elements, with greater separation between genotypes than in CPM-normalized signal.

The overlap with heterochromatin and repetitive regions is direct. BARs are overrepresented in H3K9me2/3-associated and pericentric chromatin states relative to euchromatic states, as defined by Kharchenko et al. (2011), see Figure 6B. Furthermore, the majority overlap RepeatMasker-annotated repetitive elements, and their distribution across chromosomes is correspondingly uneven (Fig. 6B). Anchor regions and H3K9me3 target regions are therefore not independent in this experiment, and normalization might reasonably be expected to remove the very signal it is meant to preserve if trimming is ineffective.

We used BASIN to evaluate previously published CUT&RUN data performed on H3K9 lysine to arginine (K9R) mutants and wild type H3K9 controls (HWT) (Stutzman et al., 2024). Remarkably, BASIN’s trimmed median approach performs robustly in this difficult context. At a pericentromeric locus on chromosome 3L, BASIN-normalized tracks recapitulate the expected loss of H3K9me3 in K9R relative to HWT, at both permissive and stringent mapping-quality thresholds (Fig. 6C). Median H3K9me3 signal in pericentric bins shows similar separation between genotypes at both MAPQ thresholds (Fig. 6D). Stratifying replicate-level effect sizes by chromatin state shows that BASIN recovers the expected genotype effects across euchromatic, Polycomb-like, heterochromatic, and pericentric compartments (Fig. 6E). The clearest discrimination between methods appears over annotated transposable elements, which are known to be marked with H3K9me3 (Fig. 6F) (Riddle et al., 2011; Sienski et al., 2012). BASIN-normalized signal shows pronounced relative enrichment of H3K9me3 in HWT compared with K9R across scaled transposable-element bodies, a separation that is greatly diminished in CPM-normalized signal at the same loci. Because CPM equalizes total signal between samples, a genotype in which a heterochromatic modification is globally reduced is rescaled upward until much of that reduction is no longer visible.

That BASIN performs well here despite substantial anchor–target overlap does not establish that the independence assumption may be disregarded. Upper-tail trimming removes anchors carrying true enrichment before the median is taken, and the result above suggests that this protection is sufficient in this dataset. In fact, approximately 1/3 of BARs in the universal panel occur within euchromatic regions, and thus the annotation-agnostic trimmed median approach allows flexibility beyond methods where reference regions are filtered based by annotation. In summary, BASIN performs remarkably well even in a dataset chosen to push its limits.

### CODA provides genotype-aware quality control of normalization output

All normalization methods, including BASIN, rest on assumptions that can fail. When further analyses and interpretation proceed from failed normalization regimes, spurious conclusions can result. Yet, removing data on nebulous grounds can also be risky. To our knowledge, normalization methods to date have not included a comprehensive framework for evaluating whether its assumptions are met and whether the resulting normalization can be trusted. With this problem in mind, we designed CODA to evaluate whether HARMONY’s assumptions are met by the input, and to evaluate the validity of normalized output at both the level of the experiment and individual samples.

CODA computes two complementary metric sets on individual samples (Fig. 7A). Baseline metrics — including BAR support, background behavior, and replicate signal structure — are computed from read-depth-normalized coverage and describe the data before BASIN is applied. Post-BASIN assessments evaluate the behavior and integrity of the normalization using the BASIN-scaled output together with HARMONY’s recorded scale factors and retained BAR sets. These signals are integrated at the experiment level, accounting for sample number, genotype-group structure, and flag severity, to yield one of four outcomes: pass, pass with caveat, manual review, or failed QC. Because the CODA framework is quite comprehensive, we provide a summary table with individual CODA metrics, where to find them, and how to interpret them (Table T2).

**Figure 7.**
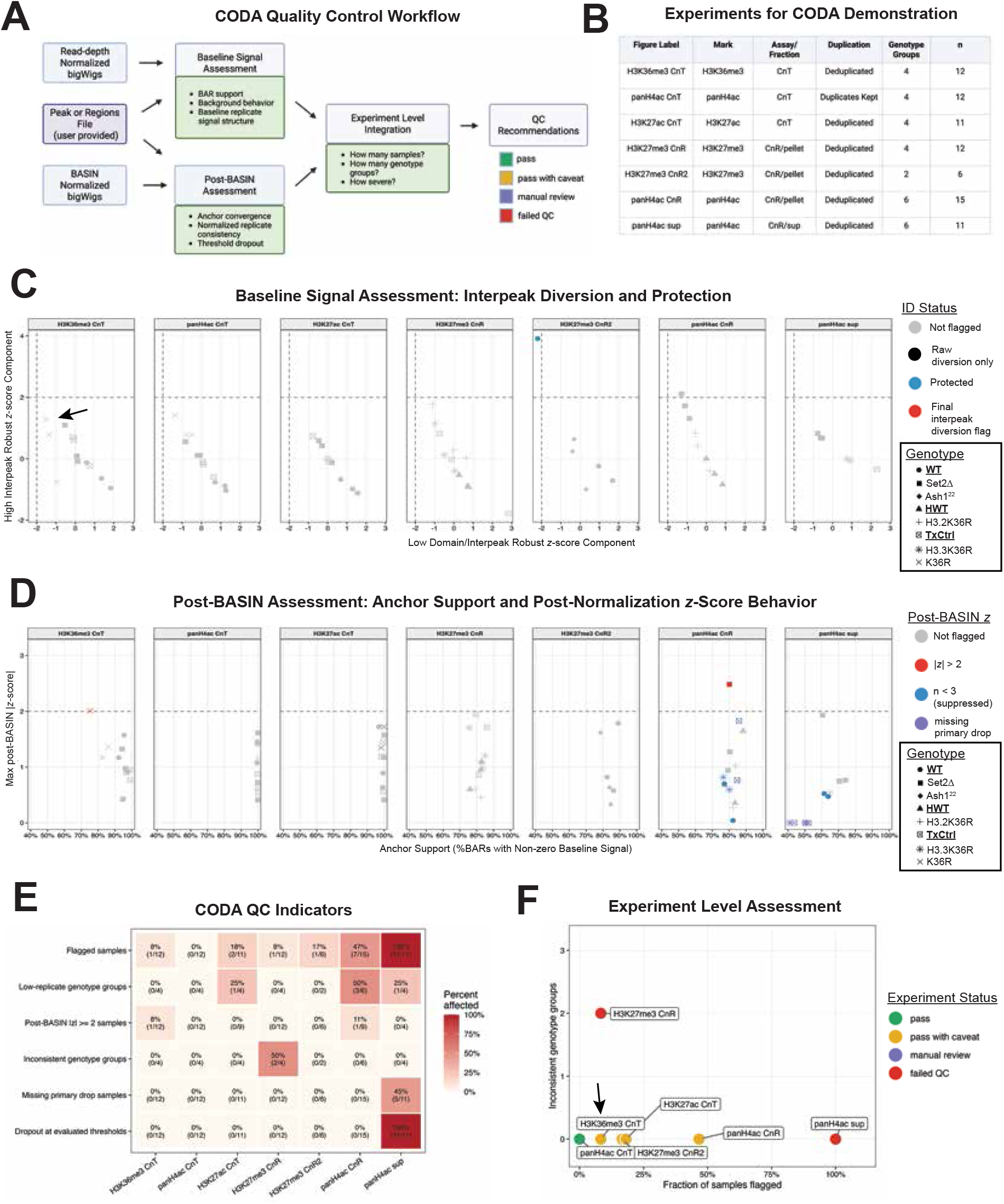
CODA provides automated, genotype-aware post-normalization quality control across diverse CUT&Tag and CUT&RUN experiments. (A) Overview of the CODA quality control workflow. Metrics capturing baseline signal behavior (BAR support, background behavior, and baseline replicate signal structure) are computed from read-depth (CPM) normalized bigWigs, while a complementary set of metrics (anchor convergence, post-normalization replicate consistency, and threshold dropout) are computed from BASIN-normalized bigWigs. A user-provided peak file that can be generated from upstream inputs is also required. These baseline and post-BASIN metrics are integrated at the experiment level — accounting for sample number, genotype group structure, and flag severity — to generate one of four experiment-level recommendations: pass, pass with caveat, manual review, or failed QC. **(B)** Summary of the seven experiments used to demonstrate CODA, spanning four histone marks (H3K36me3, panH4ac, H3K27ac, H3K27me3), two assay types (L1 CUT&Tag and wing disc CUT&RUN, the latter further split into pellet and supernatant fractions), both duplicate-retaining (panH4ac CnT) and deduplicated (all other experiments) processing, and two to six genotype groups per experiment (n = 6–15 samples per experiment). **(C)** Baseline interpeak diversion assessment for each experiment, plotted by two z-scored components of interpeak diversion (elevated interpeak signal, and reduced domain/interpeak ratio, each relative to the experiment-group median). Samples are colored by interpeak diversion status: not flagged; flagged by raw detection only; flagged but protected as likely benign given low absolute background signal; or flagged under the final QC criteria, which requires convergence of at least two of three diversion components and failure of the low-background protection clause. Point shape indicates genotype. Dashed lines mark the threshold for each component. Control genotypes are bold and underlined in the key.The black arrow indicates two H3K36R replicates that change status depending on duplicate handling (discussed in text with Fig.S6). **(D)** Post-BASIN assessment of anchor support and normalized replicate behavior for each experiment. Anchor support is the percentage of BAR regions with nonzero baseline signal for a given sample (x-axis, clipped to 40–100%); the y-axis shows the larger in magnitude of each sample’s domain or interpeak z-score, computed within its experiment group from the BASIN-scaled bigWigs at the primary drop threshold (0.20). Samples are colored by post-BASIN outlier status: not flagged; |z| > 2; z-scores suppressed because the genotype group has fewer than three replicates; or missing because no valid scale factor was computed for that sample at the primary drop threshold. Point shape indicates genotype. The dashed line marks |z| = 2. **(E)** Heatmap summarizing six sample– and group-level QC indicators across the seven demonstration experiments: fraction of samples flagged overall; fraction of genotype groups with fewer than three replicates; fraction of samples with |z| ≥ 2 post-BASIN; fraction of genotype groups showing within-genotype inconsistency; fraction of samples missing a valid scale factor at the primary drop threshold; and fraction of samples that dropped out at any evaluated drop threshold. Cell values show the affected fraction and underlying count; color indicates percent affected. **(F)** Experiment-level QC outcomes plotted against the fraction of samples flagged (x-axis) and the number of genotype groups showing within-genotype inconsistency (y-axis), colored by final CODA2 recommendation. Consistent with the CODA2 decision framework, panH4ac sup — in which all samples were flagged and every sample dropped out at one or more evaluated thresholds — receives failed QC, as does H3K27me3 CnR, which is independently flagged for having two genotype groups with within-genotype inconsistency; the remaining five experiments pass or pass with caveat. The black arrow undicates

These metrics are designed to probe different assumptions intrinsic to BASIN. Reliable HARMONY scale factors require adequate sampling of BARs. CODA therefore first measures BAR support in the baseline CPM-normalized data as the fraction and number of BAR regions with nonzero signal. Low BAR support makes the retained BAR median increasingly sensitive to sampling noise and can ultimately prevent a valid scale factor from being calculated as progressively larger fractions of high-signal BARs are removed. CODA separately records this latter behavior as threshold dropout. A second assumption is that background outside expected signal regions and BARs does not vary anomalously among otherwise comparable samples. To assess this, we developed an “interpeak diversion” metric. CODA summarizes signal over target regions, BARs, and the remaining nonrepetitive interpeak genome, and evaluates three MAD-based robust z-score components within each experiment group: elevated interpeak signal, reduced domain/interpeak signal, and elevated anchor/domain signal. Samples satisfying at least two of these three criteria are flagged for interpeak diversion. Because very low background and exceptionally tight replicate distributions can inflate relative outlier statistics, CODA suppresses the final diversion flag when group-level interpeak signal is very low and domain signal is correspondingly tight. This protection is particularly relevant to CUT&Tag datasets, which often contain substantially less interpeak signal than CUT&RUN.

CODA next evaluates whether BASIN normalization produced the expected technical behavior and whether individual samples remain coherent after scaling. Because HARMONY defines each scale factor as the reciprocal of the median signal over the retained BAR set, the median retained-BAR signal in each corresponding BASIN-normalized track should be approximately one. CODA uses the convergence of these retained-BAR medians across samples as a pipeline-integrity check: failure to converge suggests a mismatch among the BAR set, scale factor, and scaled coverage rather than a biological property of the samples. CODA also evaluates normalized domain and interpeak signal within each experiment group and flags samples that remain outliers after scaling. Additional diagnostics track dropout across trim fractions and changes in the chromosome composition of the retained BAR set, which can reveal unstable scale-factor estimation or asymmetric trimming of anchors across the genome.

CODA also evaluates the sensitivity of BASIN normalization to the BAR-trimming parameter. For each trim fraction, it measures the reproducibility of the depth-normalized medBAR across biological replicates and the relative change in HARMONY scale factors between successive trim fractions. The former identifies the trim fraction producing the most reproducible background estimate (lowest mean within-genotype medBAR CV, mean_within_genotype_medbar_cv), while the latter provides an “elbow” diagnostic showing when additional trimming produces progressively smaller changes in normalization (median_scalefactor_rel_step). CODA reports the trim fraction minimizing cross-replicate medBAR CV (suggested_drop_by_medbar_cv) as an advisory recommendation, together with the corresponding scale-factor change. Because this recommendation is based on reproducibility rather than biological ground truth, always consider the choice of drop threshold in light of the expected abundance of the target epitope and sequencing characteristics of your dataset.

We applied CODA to seven experiments spanning four histone modifications, both CUT&RUN and CUT&Tag, and two to six genotype groups per experiment (Fig. 7B). The CUT&RUN datasets included separately analyzed pellet and supernatant fractions. Baseline interpeak-diversion analysis identified samples with elevated background relative to their experiment group, together with reduced domain/interpeak signal and/or increased anchor/domain signal (Fig. 7C). Because these relative statistics can become unstable when absolute background is extremely low, CODA suppresses the final diversion call when low interpeak signal is accompanied by tight domain signal across the experiment group (Fig.7C, H3K27me3-CnR2 panel). Post-BASIN assessment then evaluates whether samples remain coherent after normalization, including retained-BAR convergence, normalized domain and interpeak behavior, and threshold dropout (Fig. 7D).

The H3K36me3 CUT&Tag experiment provides an example of a problem that depended on read processing rather than on the underlying experiment. When duplicates were retained, two K36R replicates were flagged for interpeak diversion (Fig. S6A). K36R has very little remaining specific H3K36me3 signal, making stochastic background enrichment proportionally more prominent when duplicated reads are retained. After duplicate removal, both interpeak-diversion flags disappeared (Fig. 7C). The overall fraction of flagged samples in this experiment fell from 25% to 8%, and the experiment-level assessment improved accordingly (Fig. 7E,F; FigS6C-D). Duplicate removal also produced the largest reduction in BASIN replicate dispersion for the K36R genotype in this experiment (Fig. 4E), independently supporting the interpretation that the original variability arose from duplicated background signal rather than from the normalization itself.

Across the seven experiments, QC problems were concentrated in a small number of datasets rather than distributed broadly across BASIN-normalized experiments (Fig. 7E). Two experiments met the criteria for failed QC, for different reasons. In the H3K27me3 CUT&RUN experiment, replicates were inconsistent within two genotype groups, triggering the experiment-level failure criterion for recurrent within-genotype instability. By contrast, every panH4ac supernatant sample was flagged, and scale-factor dropout occurred across one or more evaluated trim thresholds. Consequently, many samples also lacked a valid scale factor at the primary drop. The two failure modes are therefore mechanistically distinct: the H3K27me3 experiment fails because replicate behavior is internally inconsistent, whereas the panH4ac supernatant fraction fails because the BAR signal does not support stable scale-factor estimation across the trim series. Under the current decision framework, two or more inconsistent genotype groups are sufficient for failed QC, as is excessive threshold dropout.

The behavior of the panH4ac supernatant fraction is also consistent with its experimental origin. In CUT&RUN, the supernatant fraction contains released material rather than the chromatin-rich pellet, and in this dataset it did not provide a sufficiently stable BAR signal for normalization across the evaluated thresholds. Importantly, CODA does not assign this interpretation itself. It reports the observable failure — repeated loss of valid scale factors — rather than forcing a normalization value from poorly supported input. The corresponding panH4ac pellet experiment did not show this failure, illustrating that CODA evaluates the behavior of the individual experimental fraction rather than the antibody or biological comparison in isolation.

The remaining five experiments either passed or passed with caveat (Fig. 7F). Caveats reflected localized issues such as low replicate number or a small number of warning-level samples rather than broad failure of the normalization. CODA explicitly distinguishes such localized patterns from experiment-wide instability: a high fraction of flagged samples confined to one genotype group is routed to manual review, whereas the same fraction distributed across multiple genotype groups can trigger failed QC. Thus, the experiment-level recommendation retains information about both the number of affected samples and the structure of those failures across the experiment.

Together, these results show that CODA can distinguish several qualitatively different sources of concern: inadequate BAR sampling and scale-factor dropout, anomalous background behavior in individual samples, within-genotype replicate inconsistency, and mechanical failure of the scaled output to reconcile with the scale factors used to generate it. These checks are not intended to determine whether a biological result is correct. Rather, they provide an explicit framework for deciding whether BASIN’s assumptions were sufficiently supported, whether the normalization behaved as expected, and where manual inspection is warranted before quantitative interpretation.

## Discussion

BASIN addresses a central problem in quantitative chromatin profiling: preserving genuine global differences without an experimentally introduced reference. What separates it from other endogenous-reference methods is which background it uses. Other methods take their reference from enriched regions presumed unchanged or balanced, from broad bins presumed to be mostly background, or, in the case of Greenlist, from the regions most stable across IgG controls (de Mello et al., 2024). BASIN instead uses peak-like read sinks that recur in IgG controls, chosen because their share of the library does change, rising as specific signal falls. No individual anchor needs to be invariant; the method depends on the collective behavior of an empirically defined background compartment. The underlying logic is compositional: a library is a finite pool in which specific and nonspecific reads compete for representation, so each titrates the other, and anchor signal reports where that balance sits in a given sample.

What distinguishes IgG control enriched regions from diffuse interpeak background, which is also a read sink, is that it is concentrated at defined, recurrent regions. BARs therefore carry enough reads to be estimated reliably in each sample, and their recurrence across tissues and both assay chemistries makes them unlikely to reflect lineage-specific accessibility or assay-specific artifacts, whatever the source of the reads. That source is not established here: recurrent IgG signal could arise from reproducible assay– or chromatin-dependent enrichment (Jain et al., 2015; Teytelman et al., 2013), accessibility-associated untargeted cleavage or tagmentation (Hu et al., 2026; Thompson & Byrd, 2025), repeat-associated or mapping-related read accumulation (Nordin et al., 2023), or other reproducible properties that cause reads to accumulate at particular genomic regions. Anchors are not required to be biologically inert, because upper-tail trimming limits the influence of regions containing true target signal. This follows the same general robust-estimation logic as TMM and the contamination–variance tradeoff recognized by NCIS, but BASIN first restricts normalization to an empirically defined background compartment and then trims within that compartment (Liang & Keleş, 2012; Robinson & Oshlack, 2010).

Selection and trimming are both empirical and annotation-agnostic, so nothing is assumed in advance about which parts of the genome constitute background or why reads accumulate there. One consequence is that regions normally treated as problematic become useful. BARs are frequently repetitive and heterochromatic, and overlap regions excluded from locus-level interpretation because of ambiguous mapping. That caution is appropriate when assigning biological meaning to a coordinate, but it does not make the same reads uninformative about library composition. Strong, reproducible nonspecific signal in heterochromatic and blacklist-associated regions has been noted before, but has generally been filtered rather than exploited (Jain et al., 2015; Nordin et al., 2023; Teytelman et al., 2013). BASIN therefore distinguishes between reads that are unsuitable for locus-specific biological interpretation and reads that are quantitatively informative for the purpose of library scaling; the two are not necessarily equivalent.

Our validation was organized around failure modes rather than agreement with another method. A normalization can remove a real global change, create one where none exists, or be corrupted by true signal in its own reference. The H3K36me3 mutants, the cell-input titration, and the H3K9me3 dataset test these in turn, each against an independent expectation rather than an alternative normalization method. BASIN retained global changes that read-depth normalization removed, while avoiding the variability observed with the exogenous reference in the datasets examined here. These comparisons also show that replicate concordance alone cannot be used to judge normalization: a method can make replicates agree by removing biological variation that should have been retained. Because BASIN requires only the reads already in the library, it can be applied retrospectively to existing datasets, including those generated without a spike-in. Where a spike-in is present, the two methods can cross-validate each other because they fail for unrelated reasons: BASIN through background behavior or sparse anchor coverage, and spike-in through cell counting, reference addition, and recovery.

A second distinguishing feature is that the normalization model itself is evaluated rather than merely applied. OVERTURE benchmarks candidate reference panels before use, and CODA asks, for each experiment, whether BASIN’s assumptions held and whether the scale factors are technically supported. Existing QC frameworks assess library complexity, enrichment, duplication, peak structure, and replicate agreement (Boyd et al., 2021; Carroll et al., 2014; Landt et al., 2012; Zheng et al., 2020), but these do not establish that the reference behind a normalization factor behaved appropriately. CODA makes normalization failure an observable outcome and, in the datasets examined here, distinguished which assumption failed, separating inadequate anchor support, abnormal background redistribution, and replicate inconsistency. Existing tools offer a formal test of a single distributional assumption (Hicks & Irizarry, 2015), descriptive plots for the user to inspect (Angelini et al., 2015; Roule & Akizu, 2026), a single pass/fail call per sample (Jin et al., 2020), or a cross-check between two spike-ins (Patel et al., 2026), and a recent benchmark notes that it is rarely known whether a method’s assumptions are met (Ambani et al., 2026). To our knowledge, none distinguishes among multiple failure modes. We propose this finer resolution as a standard for quantitative sequencing workflows: a scale factor should be accompanied by evidence that the data support the model used to derive it.

BASIN nevertheless has important limitations. As a scalar method it cannot correct locus-specific technical bias, and it requires enough sequencing depth to sample the BAR population; very sparse background makes any background-derived scale factor unstable. The titration is also symmetric: a rise in anchor share can reflect less specific signal or more nonspecific background, and BASIN reads it as the former. A perturbation that changes nonspecific background yield across the genome, such as a global change in chromatin accessibility, would shift anchors together, and trimming, which acts on rank within a sample, cannot detect a uniform shift. Experiment-specific panels, which are more likely to retain more condition-specific accessibility regions, are the more susceptible. CODA’s interpeak-diversion metric flags atypical redistribution of background but cannot exclude every change in its composition. This vulnerability is shared by other endogenous-reference methods but not by spike-in normalization. For the same reason, BASIN-normalized values are comparable only among samples profiled with the same antibody under matched reaction conditions, not between antibodies or targets. Finally, the trimmed median should tolerate copy-number variation affecting a minority of anchors, but extensive aneuploidy or structural variation could shift a substantial fraction of the panel. Genomic DNA sequencing could be used to identify affected BARs or to make panel construction copy-number-aware, which will matter most when extending BASIN to mammalian genomes and cancer-derived cell lines.

The present implementation also uses only part of the available background. HARMONY derives its scale factor from BAR-associated signal, whereas CODA uses diffuse interpeak background only to detect abnormal redistribution. A future model could combine the two quantitatively, letting peak-like and diffuse background contribute complementary information, which could reduce sensitivity to changes in background composition. This would first require establishing how the two compartments covary across biological and technical perturbations. S3norm is a conceptual precedent in that it transforms peak and background components jointly, but its transformation assumes that common peaks and common background should be equalized between samples, which limits detection of global shifts (Xiang et al., 2020).

Although the panels developed here are specific to *Drosophila melanogaster*, the strategy should be testable elsewhere. Reproducible high-signal background loci have been described in mammalian CUT&RUN and CUT&Tag datasets (Nordin et al., 2023), and OVERTURE was designed so that panel development can proceed independently of normalization. Larger or more structurally variable genomes may need larger panels, modified recurrence criteria, or copy-number-aware filtering, though the ability to retain rather than discard repetitive sequence may be an advantage there. The same logic may extend beyond CUT&RUN and CUT&Tag: other antibody-based enrichment assays such as RIP-seq may contain analogous nonspecific populations, and unrelated assays may contain other intrinsic background whose abundance tracks library composition or recovery. A background population that is reproducible, quantitatively responsive, and empirically testable can serve as an internal reference. BASIN thus treats normalization not as an invisible preprocessing step but as a quantitative model whose reference population, assumptions, and failure modes can each be examined experimentally.

## Data and Software Availability Statement

Previously published sequencing datasets analyzed in this study are identified in Table T3. Newly generated sequencing datasets have been deposited in the NCBI Gene Expression Omnibus (GEO) and will be made publicly available upon publication of the peer-reviewed manuscript. GEO accession numbers for these datasets will be provided when the records are released. The BASIN software package, including the OVERTURE, HARMONY, and CODA pipelines, documentation, and validated *Drosophila melanogaster* BAR reference panels, is publicly available on GitHub at https://github.com/hsalzler/BASIN.

## Methods

### Study design and datasets

BASIN was developed and evaluated using IgG CUT&RUN and CUT&Tag controls, newly generated Drosophila chromatin-profiling datasets, and previously generated CUT&RUN data. A cross-experiment collection of 28 IgG libraries from eight laboratories and eight tissue or cell types was used to construct a universal background-anchor-region (BAR) panel; an additional panel was constructed from ten IgG controls associated with the L1 CUT&Tag dataset. The normalization and quality-control workflows were evaluated using L1 H3K36me3 perturbations, S2 cell-input CUT&Tag titrations, L3 wing-disc H3K9me3 CUT&RUN, and other chromatin profiles (Figs. 3–7). Dataset provenance and detailed computational procedures are provided in Supplemental Methods and Table T3.

The Set2 mutant and histone-replacement genotypes were based on previously described Drosophila systems (Larschan et al., 2007; McKay et al., 2015; Salzler et al., 2024). Histone mutants were analyzed with the appropriate matched genetic controls. Previously generated H3K27me3, H3K9me3, and pan-H4ac CUT&RUN experiments are described in their source publications (Salzler et al., 2023; Stutzman et al., 2024; Sultana et al., 2026). The pan-H4ac pellet and supernatant fractions came from the same CUT&RUN experiment and were evaluated as separate CODA groups. Ash1 mutants used for the CODA panel were *Ash1^22^/Ash1^9011^*described previously (Dorafshan 2019) alongside WT controls.

### Western blotting and densitometry quantification

For each sample, 50 L1 larvae were crushed with micropestles in 100 μL of working lysis buffer (LB) solution per tube. The lysis buffer solution contained 83 μL of SUTEB (1% SDS, 8M urea, 10 mM Tris-Cl, pH 8.0, 10 mM EDTA, pH 8.0), 5 μL of 5% β-mercaptoethanol (BME), 2 μL 2.55 M Sodium Butyrate, and 10 μL of 100x protease inhibitor cocktail (Thermo Fisher Scientific, #78429). Samples were sonicated via Bioruptor Pico (Diagenode) for 10 cycles, 30 seconds on, 30 seconds off. Lanes were loaded with 40ug of total protein and run on 4-15% Mini-PROTEAN TGX Stain-Free protein gels (Bio-Rad, #4568086). The gel was transferred to a nitrocellulose membrane (Bio-Rad #170-4270) at 25 V (1.3 A) for 7 minutes via the Bio-Rad Trans-Blot Turbo transfer system (Bio-Rad #100-240VAC). Blots were incubated overnight at 4 °C with a primary antibody, e.g. H3K36me3 (abcam #ab9050, 1:2000) or H4 (GenuIN Biotech (#2134, 1:5000) in TBS-T, then incubated with Anti-rabbitIgG-HRP conjugate (Sigma #12-348, 1:5000) dilution for 1 hour at RT. H3K36me3 and H4 probing were performed serially without stripping to prevent depletion of H4 signal. Chemiluminescence (Bio-Rad #34580) was performed and imaging on Amersham Imager 680 (GE Healthcare) prior to densitometry quantification. Separate gels from the same lysates were run and stained with Colloidal Blue (Invitrogen #LC6025) total protein stain to provide an alternative normalization reference.

Western blot images were saved as.tif files and quantified using ImageJ software (Schneider et al., 2012). Band intensities were measured using background-subtracted integrated densities. For all analyses, H3K36me3 was normalized to its corresponding H4 (Fig. 4A) or total protein (Fig. S3) control intensity within each lane. Geometric means and 95% confidence intervals for 5 replicate pairs per comparison were displayed. Statistical significance was assessed using two-tailed paired t-tests on log2-fold change values (p < 0.05 (*), p < 0.01 (**), and ns (not significant), Fig. 4A.

### Chromatin profiling, sequencing, and alignment

New CUT&Tag experiments followed published CUT&Tag procedures (Kaya-Okur et al., 2019), with assay-specific preparation, sequencing, and alignment details described in Supplemental Methods. The newly generated Ash1 CUT&RUN dataset used previously published experimental and sequencing-processing procedures (Salzler et al., 2023). Previously generated CUT&RUN data were reanalyzed from aligned BAMs; their wet-bench and original alignment methods are cited to the relevant publications. Public IgG raw reads used for BAR discovery were realigned uniformly to dm6, whereas three supplied L1 IgG BAMs were used as aligned inputs. All downstream genomic coordinates refer to the Drosophila melanogaster dm6 assembly.

### OVERTURE: discovery and selection of background anchor regions

OVERTURE identifies recurrent peak-like read-enrichment regions in IgG controls rather than requiring individual invariant genomic loci. For the Drosophila universal panel, 1-bp CPM coverage tracks were generated from prepared IgG BAMs and subjected to SEACR peak calling under alternative thresholds and stringency settings (Meers et al., 2019a). Regions supported by prespecified fractions of IgG samples were assembled into consensus candidate BAR panels, with short intervals optionally excluded. The selected universal panel was derived from relaxed-mode SEACR peaks at threshold 0.005, with 90% recurrence and exclusion of intervals shorter than 50 bp.

Candidate panels were assessed using six L1 CUT&Tag targets and matched biological replicates, examining BAR support, within-genotype variability and genotype shifts (Fig. 3; Fig. S2). The validated universal Drosophila BAR panel is supplied with the software. The OVERTURE workflow permits construction and evaluation of candidate panels from IgG controls in other experimental systems. Specific candidate grids and panel-selection metrics are described in Supplemental Methods.

### HARMONY: background-anchored normalization

HARMONY calculates a sample-specific scale factor from coverage over a fixed BAR panel. Using matched processing settings, it measures per-BAR raw signal, removes regions with zero signal in every analyzed sample, and ranks the remaining BAR values within each sample. The highest-signal fraction d is discarded, and the median signal over the retained BARs (medBAR) defines the scale factor as 1/medBAR. The default primary drop fraction was 0.20; additional fractions were evaluated for sensitivity. Each sample is scaled independently, without imposing equal target-enriched signal across genotypes.

CPM– and BASIN-normalized bigWigs were generated using deepTools bamCoverage (Ramírez et al., 2016). BASIN tracks were generated by applying the calculated sample-specific scaling factors directly during coverage generation. Corresponding CPM tracks were generated using matched read-processing and coverage settings. HARMONY records scale factors and retained regions for downstream quality control; implementation details and handling of invalid scale factors are given in Supplemental Methods.

### Validation analyses

For L1 H3K36me3 validation, the primary analysis used duplicate-removed libraries owing to high duplication, with duplicates-retained and E. coli spike-in comparisons analyzed separately (Fig. 4; Figs. S4–S5). BARs intersecting the copy-number-variable transgenic histone array were removed from the relevant panels before scaling. Gene-body H3K36me3 signal was measured over dm6 intervals derived from FlyBase annotation release 6.38 using deepTools multiBigwigSummary. After collapsing identical genomic intervals, 16,621 unique intervals remained. The highest-ranked 25% (4,155 intervals) were selected using median CPM-normalized H3K36me3 signal across three WT control replicates; this same set was applied to every genotype and normalization method. All-interval summaries were also examined. Mutant effects were expressed relative to the median of the appropriate matched-control replicates; detailed calculations are in Supplemental Methods.

For S2 CUT&Tag titrations, H3K4me1 and H3K27me3 signals were compared across eight cell inputs with a fixed 200-K562-cell reference per reaction (Fig. 5). CPM, BASIN, and K562-reference normalization were compared using fixed genomic interval sets; signal-versus-cell-input trends were evaluated against expected log-log slopes as detailed in Supplemental Methods.

For the L3 wing-disc H3K9me3 analysis, three HWT and three K9R biological replicates were compared using independently supplied MAPQ ≥0 and MAPQ ≥10 BAM sets (Fig. 6). BASIN did not impose an additional MAPQ filter on these inputs. Signal was assessed over Kharchenko-derived chromatin-state intervals and pericentric extensions, and scaled profiles were generated across annotated transposable elements. The universal BAR panel was characterized separately for chromatin-state assignment and repeat overlap. Interval definitions and profile calculations are described in Supplemental Methods.

### CODA: integrated quality-control analysis

CODA examines whether background scaling behaves as expected, using matched CPM and BASIN tracks, retained BARs, target-enriched intervals and background interpeak regions. Seven QC groups derived from five profiling datasets were evaluated, including separate pan-H4ac pellet and supernatant groups (Fig. 7). Diagnostics encompass BAR support, relative enrichment of target, BAR and interpeak compartments, within-genotype reproducibility, drop-fraction stability, and post-normalization outliers. Sample– and genotype-level observations are integrated into experiment-level classifications (pass, pass with caveat, manual review or fail), using the prespecified rules in the distributed CODA implementation. Thresholds, safeguards and figure-specific computations are provided in Supplemental Methods.

## Supplemental Figures and Tables

**Figure S1.**
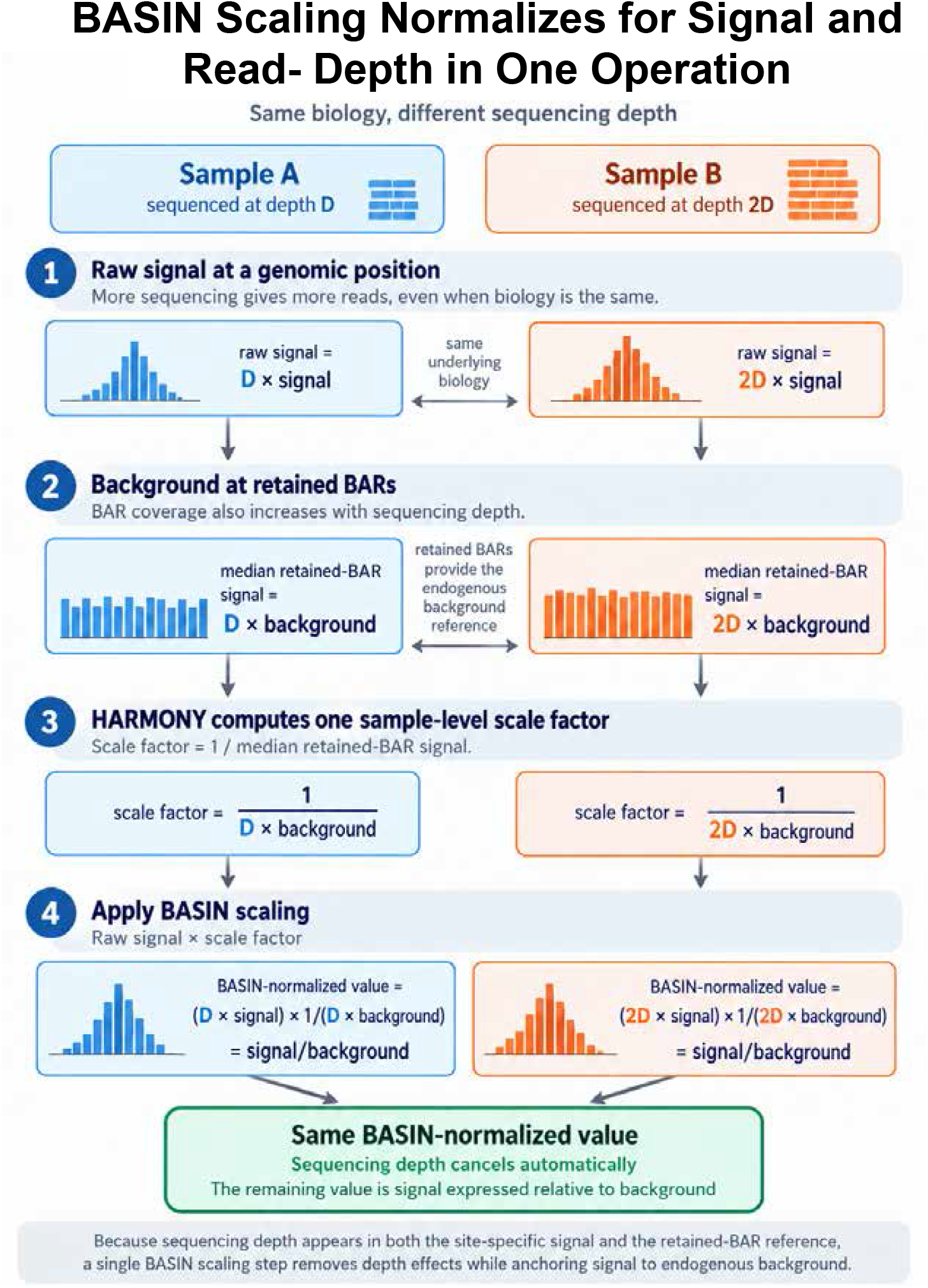
A single BASIN scaling step removes sequencing-depth differences and anchors signal to background. Schematic of two samples with the same underlying biology sequenced at depths D (Sample A) and 2D (Sample B). **(1)** Raw signal at a genomic position scales with sequencing depth. **(2)** Median coverage over the retained BARs scales with depth in the same way. **(3)** HARMONY computes one scale factor per sample, 1/(median retained-BAR signal). **(4)** Multiplying raw signal by the scale factor cancels the depth term, so both samples yield the same BASIN-normalized value, expressed as signal relative to background.

**Figure S2.**
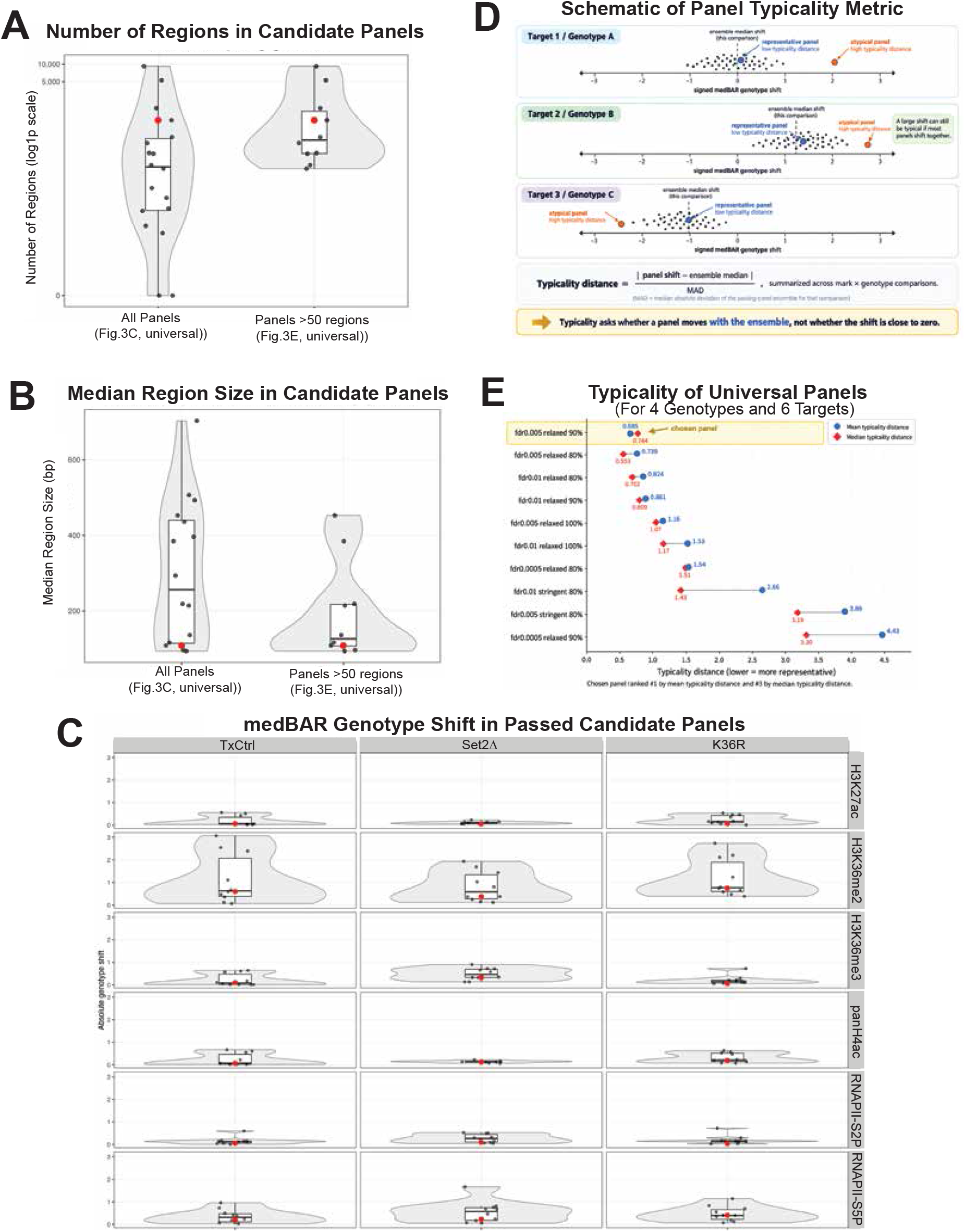
Properties of candidate universal BAR panels and genotype-shift typicality of the selected panel. Supplementary Figure S2. Properties of candidate universal BAR panels and genotype-shift typicality of the selected panel. (A) Number of regions in all candidate universal panels (with regions <50bp filtered out, Fig. 3C) (left) and in panels with more than 50 regions (from Fig. 3E) (right), on a log1p scale. Each point is one parameter combination and the red point is the selected panel; violins show the distribution and boxes the median and interquartile range. **(B)** Median region size (bp) for the same two sets of panels, plotted as in (A). **(C)** Absolute medBAR genotype shift across passing panels for each test genotype (TxCtrl, Set2Δ, K36R) and each target in the L1 CUT&Tag evaluation dataset (rows: H3K27ac, H3K36me2, H3K36me3, pan-H4ac, RNAPII-S2P, and RNAPII-S5P). Shifts are relative to the reference genotype. The red point is the selected panel. **(D)** Schematic of the typicality metric. For each target-by-genotype comparison, a panel’s signed medBAR genotype shift is compared with the median shift of the panel ensemble from Fig. 3E, and divided by the median absolute deviation (MAD) of that ensemble. A panel whose shift lies close to the ensemble median has a low typicality distance even when the shift itself is large. **(E)** Mean (blue circles) and median (red diamonds) typicality distance across all target-by-genotype comparisons for the ten passing universal panels, ordered by mean distance. The selected panel (SEACR threshold 0.005, relaxed, 90% consensus; highlighted) ranked first by mean and third by median typicality distance.

**Figure S3.**
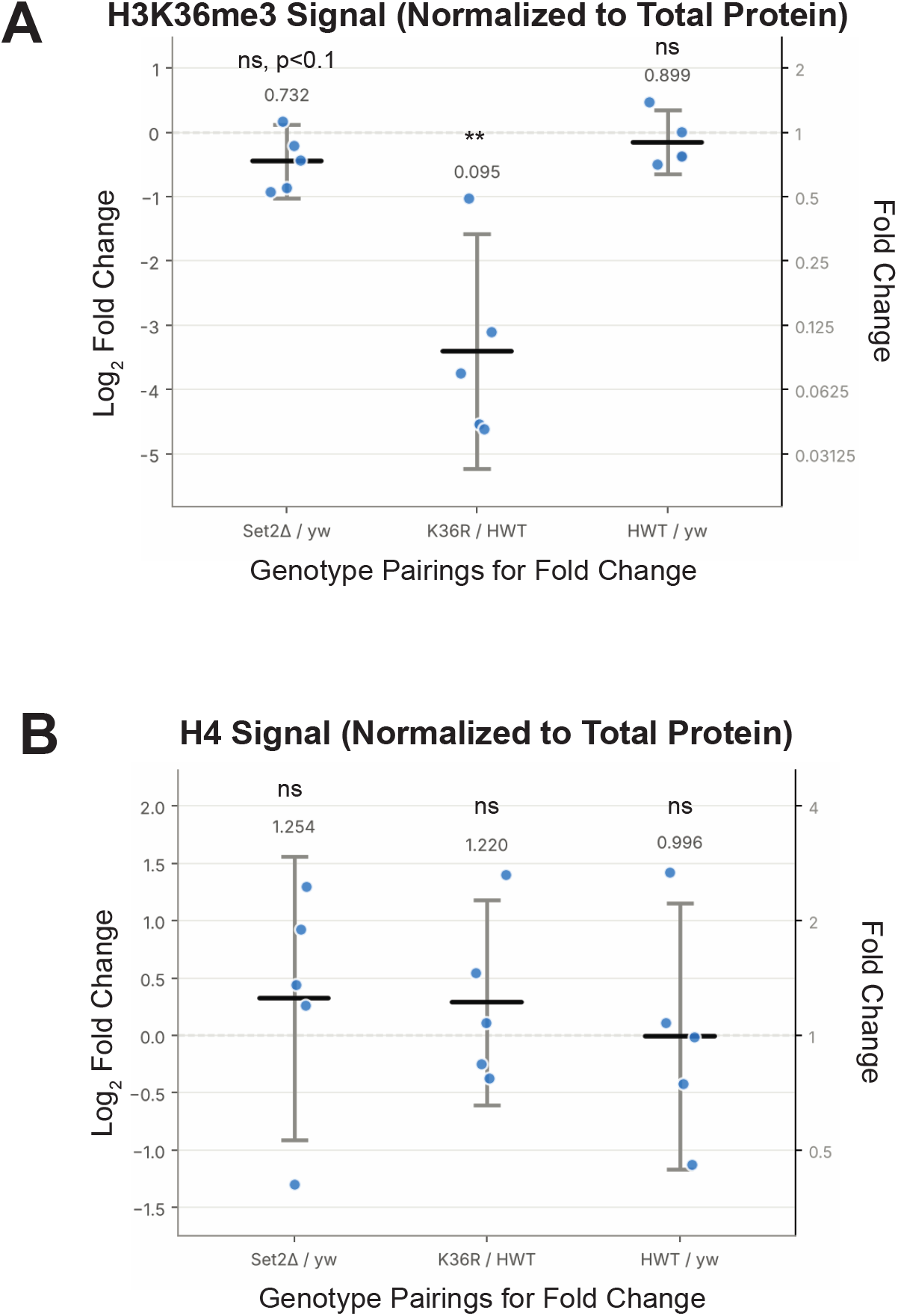
Total-protein normalization of H3K36me3 western blots and assessment of H4 as a loading control. **(A)** Quantification of H3K36me3 western blots normalized to total protein for 5 biological replicates per genotype. H3K36me3 band intensities were normalized to colloidal blue total protein stain of a gel loaded in parallel from the same lysates. Normalized intensities were expressed as both fold changes and log2 fold changes relative to the corresponding matched control for each biological replicate. The geometric mean (GM) of each comparison (mutant/control or control/control) is reported above each column, along with statistical significance. Points represent individual biological replicates; horizontal bars indicate the mean log2 fold change, and error bars indicate the 95% confidence interval. Statistical significance was calculated using two-tailed paired t-tests on log2 fold-change values (**p < 0.01; ns, not significant; ns, p < 0.1 indicates 0.05 ≤ p < 0.1). **(B)** H4 band intensities from the same blots, normalized to total protein and plotted and tested as in (A). No significant difference in H4 relative to total protein was detected for any comparison, consistent with the use of pan-H4 as a loading control in Figure 4A.

**Figure S4.**
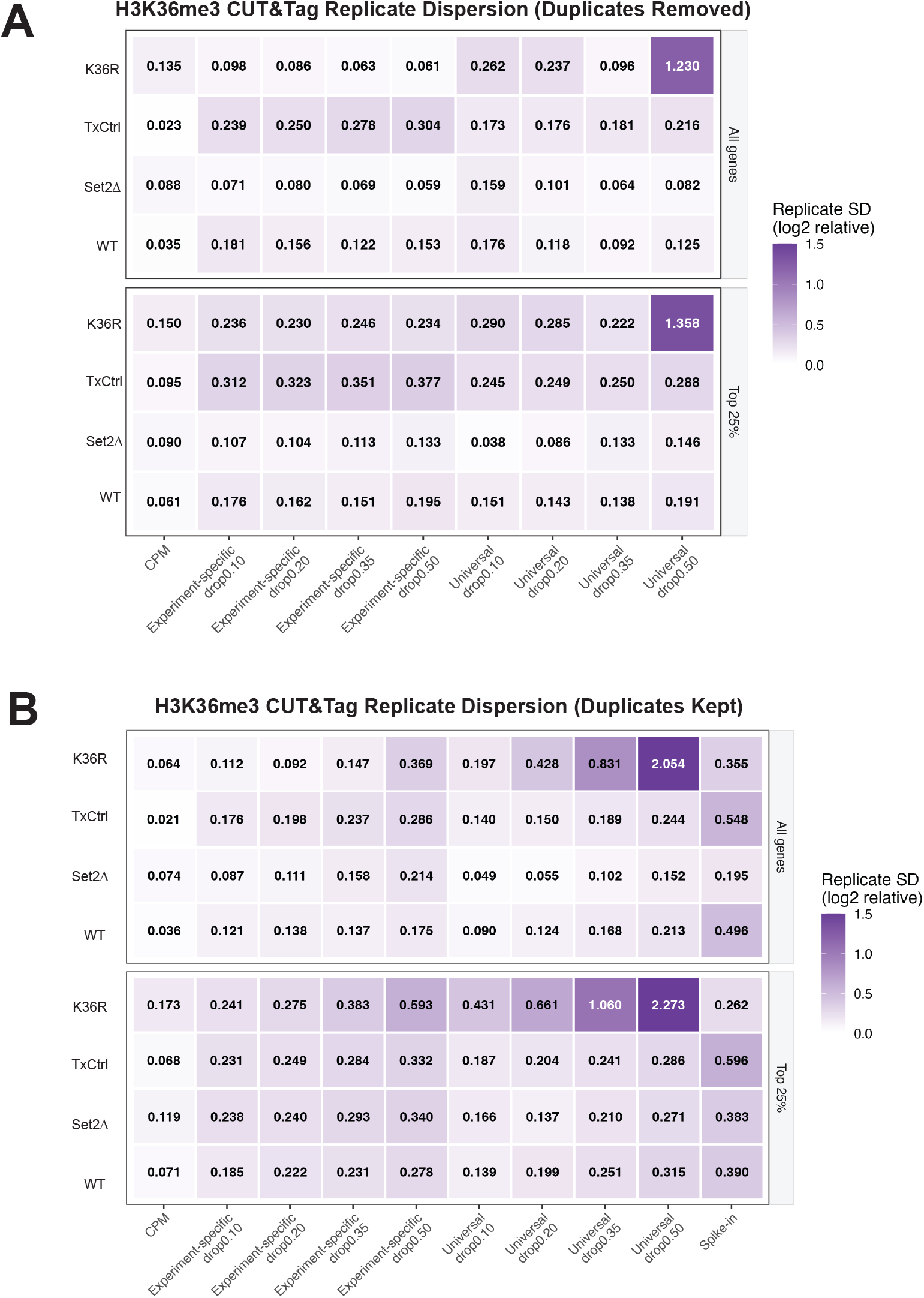
Replicate dispersion of H3K36me3 signal across BAR panels and anchor-drop thresholds. **(A)** Replicate dispersion after duplicate removal for CPM and for BASIN with the experiment-specific or universal panel at drop thresholds of 0.10, 0.20, 0.35, and 0.50, shown for all genes (top) and the top quartile of H3K36me3-marked genes (bottom). Replicate dispersion was calculated as in Figure 4E; cell values give the standard deviation. **(B)** The same analysis with duplicate reads retained, with spike-in normalization (*E. coli* carry-over reads) added for comparison. The color scale is shared between (A) and (B) and saturates at 1.5.

**Figure S5.**
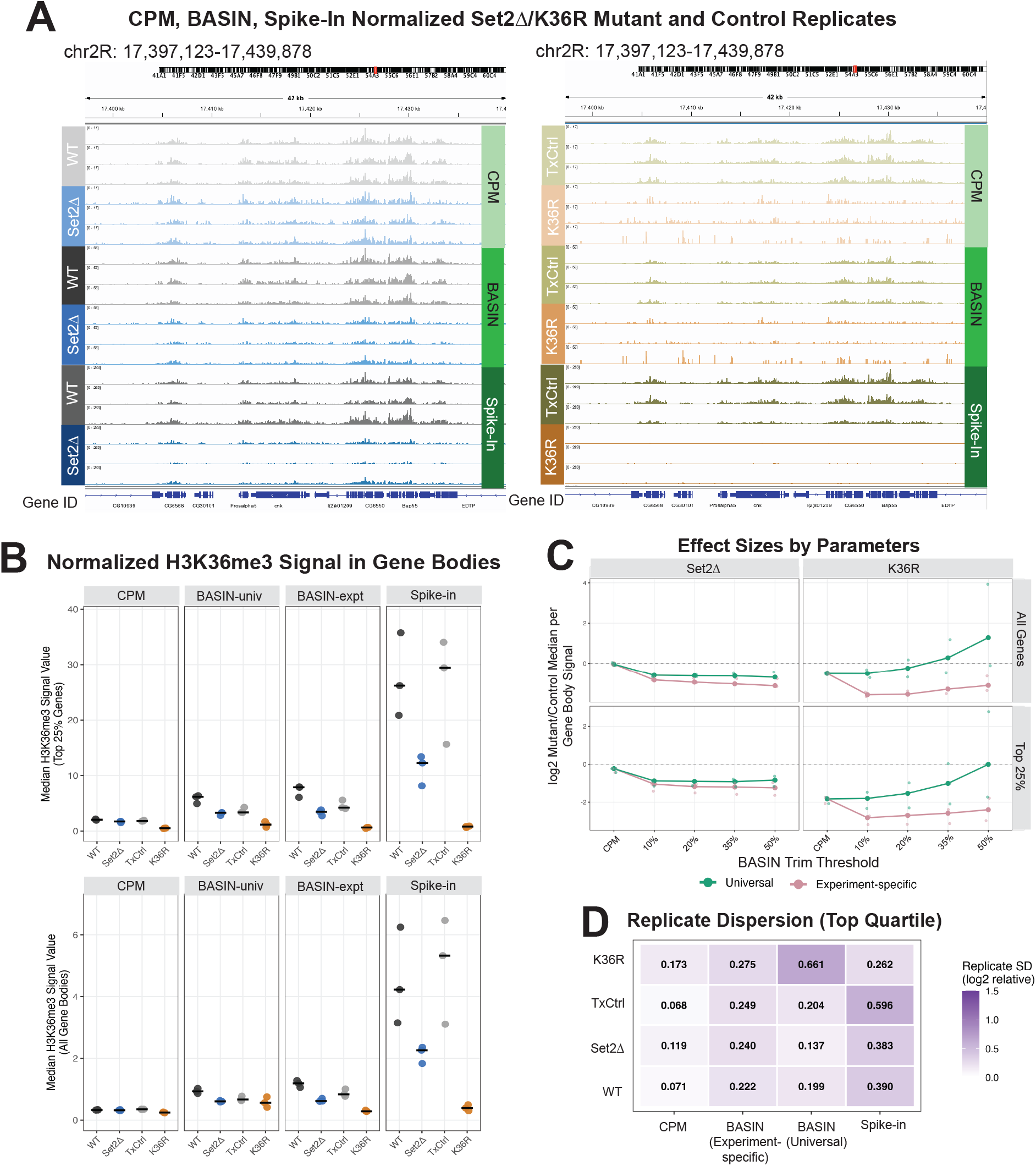
BASIN normalization detects expected H3K36me3 depletion in Set2Δ and H3K36R mutants using duplicate-retaining alignments. **(A)** Representative genome-browser views (chr2R:17,397,123–17,439,878) of H3K36me3 profiles for Set2Δ and H3K36R mutant replicates and their matched controls, following CPM, BASIN (drop0.20), or spike-in normalization. Tracks are displayed on identical scales within each normalization method. Spike-in normalization used *E. coli* carry-over reads. **(B)** Median normalized H3K36me3 signal across gene bodies of genes with the top quartile of H3K36me3 signal for each normalization method. Each point represents one biological replicate; horizontal bars indicate the median of three replicates. Gene-body signal was quantified using the same summary metric across all normalization methods; *normalization units are not comparable between methods.* **(C)** Relative effect sizes for Set2Δ and H3K36R mutants across BASIN anchor-drop thresholds, for all genes (top) and the top quartile of H3K36me3-marked genes (bottom). Values represent the median log₂ ratio of mutant signal relative to the median of the corresponding control genotype. Universal and experiment-specific anchor strategies produce similar estimates for Set2Δ and diverge for K36R, where universal-panel estimates rise toward and above zero at higher thresholds. **(D)** Replicate dispersion for the top quartile of H3K36me3-marked genes, calculated as in Figure 4E, for CPM, BASIN (drop0.20) with the experiment-specific or universal panel, and spike-in normalization. Lower values indicate greater agreement among biological replicates.

**Figure S6.**
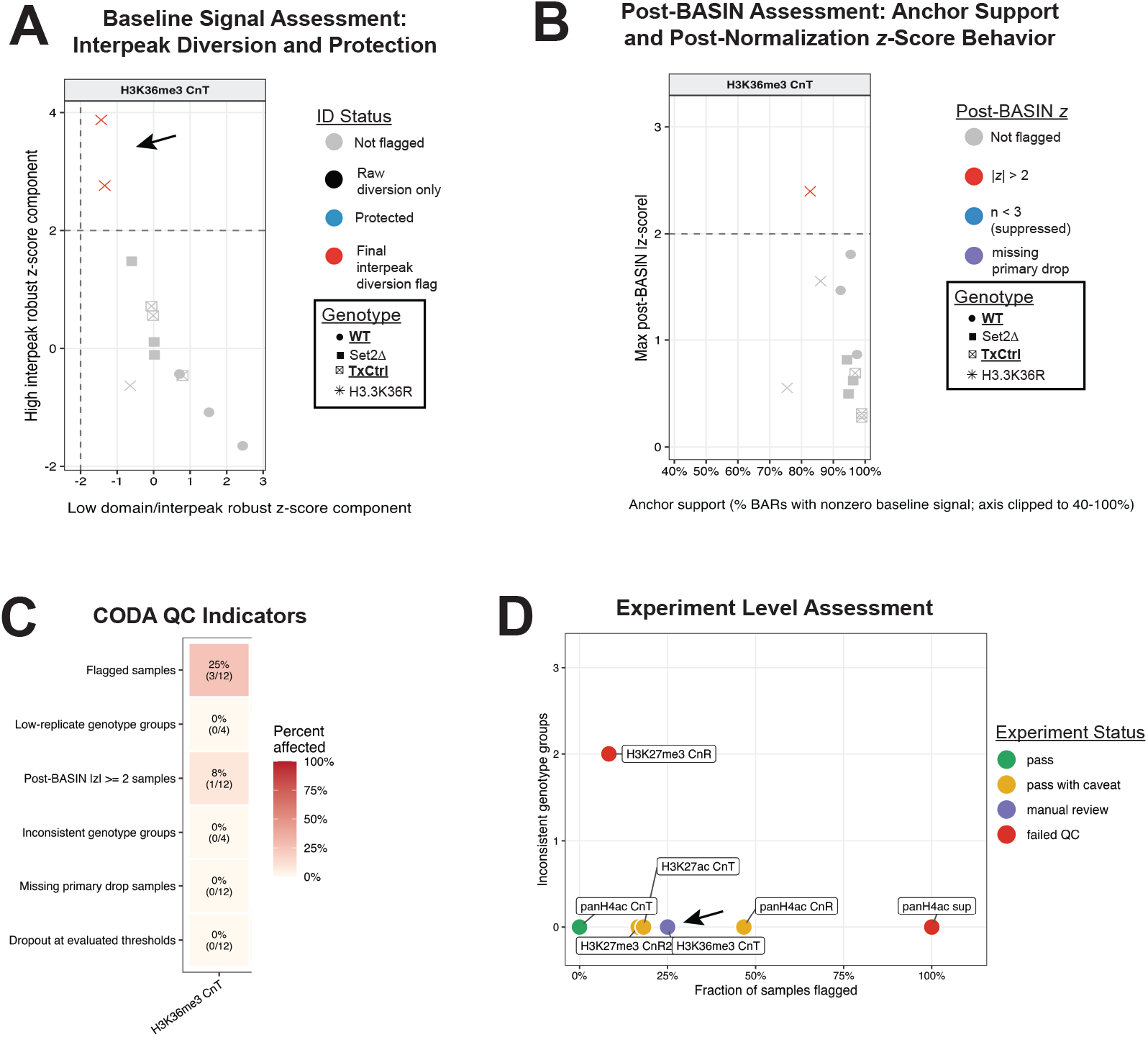
CODA assessment with duplicate reads retained in the H3K36me3 CUT&Tag experiment. CODA output for the H3K36me3 CUT&TAG experiment in Figure 7, with duplicate reads retained. **(A)** Baseline interpeak-diversion assessment, plotted as in Figure 7C. Two K36R replicates in the H3K36me3 CUT&Tag experiment receive the final interpeak-diversion flag (black arrow). **(B)** Post-BASIN assessment of anchor support and normalized replicate behavior, plotted as in Figure 7D. **(C)** CODA QC indicators, as in Figure 7E. With duplicates retained, 25% (3/12) of H3K36me3 CUT&Tag samples are flagged, compared with 8% (1/12) after duplicate removal. **(D)** Experiment-level assessment, as in Figure 7F (all experiments included). With duplicates retained, the H3K36me3 CUT&Tag experiment is assigned manual review (black arrow).

## Supplemental Tables

**Table T1.** Library QC. Statistics for L1 and titration CUT&Tag experiments. (A) L1 larval CUT&Tag. There are 79 libraries across seven targets: H3K27ac, H3K36me2, H3K36me3, panH4ac, RNAPII-S2P, RNAPII-S5P, and IgG. Four genotypes were profiled: WT, TxCtrl, *Set2*Δ, and *H3K36R*. Each library comes from 50,000 lightly fixed with 0.1% formaldehyde for 1 min and tagmented with pAG-Tn5. Tagmentation conditions differed by target: CUTAC-DMF buffer for H3K27ac, RNAPII, and IgG; 300 mM Mg²⁺ buffer for H3K36me2/3 and panH4ac. Columns list experiment number, scale, fixation, primary and secondary antibodies, Tn5, tagmentation buffer, genotype, target, and i7/i5 index adapters. Read accounting gives raw reads, reads after adapter clipping, reads mapped to dm6, and reads mapped to *E. coli*. *E. coli* reads come from carryover DNA in the pAG-Tn5 preparation, not from an added spike-in. Spike-in scale factors were calculated as 10,000 / *E. coli* read count, and normalized coverage is dm6 reads × scale factor; both are shown for comparison only (see main text). The remaining columns give the *E. coli*:dm6 read ratio, the fractions of reads mapping to dm6, *E. coli*, and chrM, the duplicate fraction, f(dup), and estimated library size. **(B) S2 cell titration CUT&Tag.** There are 16 libraries: H3K4me1 and H3K27me3, each at eight S2 cell inputs from 100,000 down to 750 cells (a 133-fold range). Each S2 sample was mixed with 200 human K562 nuclei as an exogenous spike-in. Tagmentation used pAG-Tn5 (1:20) in 300 mM Mg²⁺ buffer at 25 °C. Read accounting gives raw reads, dm6-mapped reads, human/mouse-mapped reads, and unassigned reads, plus the fraction of each and the chrM fraction. See supplemental data file.

**Table T2.** CODA metric reference. Metrics and diagnostics reported by CODA, grouped as essential (inspect for every run), frequently useful (common diagnostics), or other/advisory (supporting detail, parameter sensitivity, or deep-dive outputs). For each metric the table gives the stage and level at which it is computed, what it means, the output file and column(s) in which it is reported, and the rule or interpretation applied. Thresholds are the CODA2 defaults. See supplemental data file.

**Table T3.** Datasets used in this study. Public and newly generated *Drosophila melanogaster* CUT&RUN (CnR) and CUT&Tag (CnT) datasets used for BAR panel construction, BASIN validation and CODA evaluation. Twenty-eight IgG control samples from 12 GEO series (10 public, 2 from this study) were used to build the universal BAR panel. Experiment-specific IgG panels were built from genotype-matched L1 IgG controls. Antibody-targeted datasets were used for panel selection, titration, H3K36me3 validation, H3K9me3 validation, and CODA analysis, as indicated. Description gives tissue or cell type, assay, and target. Purpose gives sample counts, number of genotypes, and the analysis each dataset was used for. Datasets are listed in order of appearance. Citation gives the PubMed ID of the original publication. Individual GSM accessions for samples used from public series are listed in Table S1. S2, *Drosophila* Schneider 2 cells; L1, first-instar larvae; L3, third-instar larvae; NC14, nuclear cycle 14; RNAPII-S2P and – S5P, RNA polymerase II phosphorylated at Ser2 and Ser5 of the CTD.

| GEO | Description | Antibodies (new studies) | Purpose | Figure | Citation |
|---|---|---|---|---|---|
| GSE195654 | S2 cells, CnT | NA | 4 IgG samples, universal panel | 3 | PMID: 36192462 |
| GSE270072 | L3 brain, CnT | NA | 3 IgG samples, universal panel | 3 | PMID: 40855028 |
| GSE306499 | Adult Head, CnT | NA | 1 IgG sample, universal panel | 3 | PMID: 41066503 |
| GSE263999 | Embryo, NC14, CnR | NA | 1 IgG sample, universal panel | 3 | PMID: 40855028 |
| GSE247152 | S2 cells, CnR | NA | 3 IgG samples, universal panel | 3 | PMID: 40527990 |
| GSE247151 | S2 cells, CnR | NA | 2 IgG samples, universal panel | 3 | PMID: 40527990 |
| GSE247377 | L3 leg disc, CnR | NA | 2 IgG samples, universal panel | 3 | PMID: 42206359 |
| GSE211269 | L3 wing disc, CnR | NA | 2 IgG samples, universal panel | 3 | PMID: 37211836 |
| GSE255491 | L3 wing disc, CnR | NA | 2 IgG samples, universal panel | 3 | PMID: 40063800 |
| GSE247150 | Adult ovary, CnR | NA | 3 IgG samples, universal panel | 3 | PMID: 40527990 |
| Pending public release | L3 wing disc, CnR, supernatant and pellet separately | Sigma 269A-17 | 2 IgG samples, universal panel | 3 | This Study |
| Pending public release | L1 CnT, IgG | Abcam ab46540 | 3 WT IgG samples, universal panel | 3 | This Study |
| Pending public release | L1 CnT, IgG | Abcam ab46540 | 10 samples, 4 different genotypes, experiment specific panel | 3,4 | This Study |
| Pending public release | L1 CnT, H3K36me3 | Epicypher 13-0031 | 12 samples, 4 genotypes, panel selection, BASIN validation, CODA | 3, 4, 7,S2, S3, S4 | This Study |
| Pending public release | L1 CnT, panH4ac | Active Motif 39925 | 12 samples, 4 genotypes, panel selection, CODA | 3, 7, S2 | This Study |
| Pending public release | L1 CnT, H3K27ac | Epicypher 130059 | 11 samples, 4 genotypes,panel selection, CODA | 3, 7, S2 | This Study |
| Pending public release | L1 CnT, H3K36me2 | Epicypher 130056 | 12 samples, 4 genotypes, panel selection | 3, S2 | This Study |
| Pending public release | L1 CnT, RNAPII-S2P | CST #E1Z3G | 11 samples, 4 genotypes,panel selection | 3, S2 | This Study |
| Pending public release | L1 CnT, RNAPII-S5P | CST #D9N5I | 11 samples, 4 genotypes,panel selection | 3, S2 | This Study |
| Pending public release | S2 cells, CnT, H3K4me1 | Epicypher 130057 | 8 samples, titration vaidation | 5 | This Study |
| Pending public release | S2 cells, CnT, H3K27me3 | CST #9733 | 8 samples, titration validation | 5 | This Study |
| GSE236964 | L3 wing disc, CnR, H3K9me3 | NA | 6 samples, 2 genotypes, K9me3 validation | 6 | PMID: 38795706 |
| GSE215017 | L3 wing disc, CnR, H3K27me3 | NA | 12 samples, 4 genotypes, CODA | 7 | PMID: 36857457 |
| Pending public release | L3 wing disc, panH4ac CnR, pellet | NA | 15 samples, 6 genotypes, CODA | 7 | PMID: 42990221 |
| Pending public release | L3 wing disc, panH4ac CnR, supernatant | Active Motif 39925 | 11 samples, 5 genotypes | 7 | This Study |
| Pending public release | L3 wing disc, H3K27me3, CnR | CST #9733 | 2 genotypes, 6 samples | 7 | This Study |

